# Sleep Disruption Improves Performance in Simple Olfactory and Visual Decision-Making Tasks

**DOI:** 10.1101/2024.11.02.621641

**Authors:** Paula Pflitsch, Nadine Oury, Kumaresh Krishnan, William Joo, Declan G. Lyons, Maxim Capelle, Kristian J. Herrera, Armin Bahl, Jason Rihel, Florian Engert, Hanna Zwaka

## Abstract

Sleep disruption drastically impacts cognitive functions including decision-making and attention across many different species. In this study, we leveraged the small size and conserved vertebrate brain structure of larval zebrafish to investigate how sleep disruption modulates visual-and olfactory-decision-making. Strikingly, sleep disruption improved performance in both paradigms. Specifically, sleep disruption lengthens reaction times and increases correct decisions in a visual motion discrimination task, an effect that we attribute to longer integration periods in disrupted animals. Using a drift diffusion model, we predict specific circuit changes underlying these effects. Additionally, we demonstrate that sleep disruption heightens odor sensitivity in an olfactory decision-making task, likely mediated by cortisol. Our findings lay essential groundwork for investigating the brain circuit changes that arise from sleep disruption across species.

## INTRODUCTION

We spend about one-third of our lives sleeping, yet the functions and mechanisms of sleep are still not fully understood. The cognitive impairments caused by lack of sleep, however, underline that sleep is essential for normal brain function. Disruption of sleep affects numerous aspects of our body and brain, including attention, memory, and decision-making ^1–7^. Yet, the precise mechanisms by which cognitive function is compromised after sleep disturbance remain unclear. To understand how sleep loss affects cognitive functions like decision-making, we turn to larval zebrafish (Danio rerio), which display a robust sleep-wake cycle similar to humans ^8^ and are well-suited to high-throughput experiments due to their small size and rapid maturation. Zebrafish also share conserved brain architecture and neurochemical pathways with mammals, including the hypocretin ^8^, melatonin ^9,10^, galanin ^11^, serotonergic raphe ^12^ and cortisol ^13,14^ systems, suggesting a high degree of conservation in sleep mechanisms ^15,16^.

Recently, the concept that sleep homeostasis can be measured solely by delta power has been challenged ^17^. After sleep deprivation, slow-wave brain activity initially increases but returns to normal relatively quickly, whereas behavioral changes persist for a longer period, indicating a slower homeostatic recovery in behavior. This highlights the importance of examining how sleep disruption affects behavior. The larval zebrafish offers a unique opportunity to examine the effects of sleep disruption at both behavioral and molecular levels. Various methods exist for perturbing sleep in larval zebrafish, including mechanical disturbance ^9,18–20^, water flow ^21^, acoustic stimulation ^18,22^, optic stimulation ^11^, or light exposure ^8,18,23,24^. Sleep in larval zebrafish is classically defined as behavioral quiescence lasting at least one minute, which correlates with increased arousal threshold and is homeostatically regulated ^8,9,22,25,26^.

Larval zebrafish exhibit well-studied decision-making behaviors with known circuit models ^27–29^. One of these is the optomotor response (OMR) ^27,30,31^, an innate behavior common to fish and insects ^30,32^, where animals stabilize themselves in response to whole-field visual motion, for example in water currents. This behavior can be reproduced in a simple decision-making task using a moving dot stimulus ^27^, with performance easily quantified through discrete swimming events (swim bouts). The hindbrain neuronal circuits underlying OMR are well characterized ^27,33^, allowing us to investigate how sleep disruption alters specific circuitry to modulate behavior. Another important decision-making behavior in larval zebrafish is avoidance of stimuli that could signal danger. Larvae use olfactory cues to assess potential threats in their environment, leading to avoidance behavior ^34,35^. Zebrafish possess high-affinity olfactory receptors for scents emitted by decaying flesh, like cadaverine and putrescine, which elicit strong aversive responses ^36,37^. Thus, larval zebrafish provide a model for studying the consequences of sleep disruption on perceptual decision-making across multiple sensory modalities.

In this study, we analyzed the effects of sleep disruption on both visual-based and olfactory-based decision-making in larval zebrafish. We first established that light can be used to perform high-throughput sleep disruption experiments in larvae. We then assessed performance in an optomotor task ^27^ before and after sleep disruption, revealing that sleep disturbance enhanced optomotor performance through a speed-accuracy trade-off, an effect that could be replicated with melatonin treatment. In contrast, in an olfactory task, sleep disruption increased odor avoidance, likely mediated by cortisol, which prompted earlier turning away from the odor source.

## RESULTS

### Larval zebrafish sleep can be sleep disrupted using light

To investigate how sleep disruption impacts perceptual decision-making in larval zebrafish, we developed a light-based sleep disruption protocol. Using visible light, we introduced different light intervals during both day and night: Constant light (Fig. 1), light pulses, and 6 h of premature light during the night (Fig. S1). Control groups were kept under a standard 14 h light /10 h dark cycle. Sleep was quantified using automated movement tracking with a high-speed camera, defining sleep bouts as periods of inactivity lasting 1 minute or more (Fig. 1a, b). We found that light exposure drastically reduced nighttime sleep compared to unperturbed controls (Fig. 1b-e, S1a-d). Additionally, sleep-deprived fish exhibited increased sleep the following morning, indicating a rebound effect (Fig. 1d, e).

**Figure 1:**
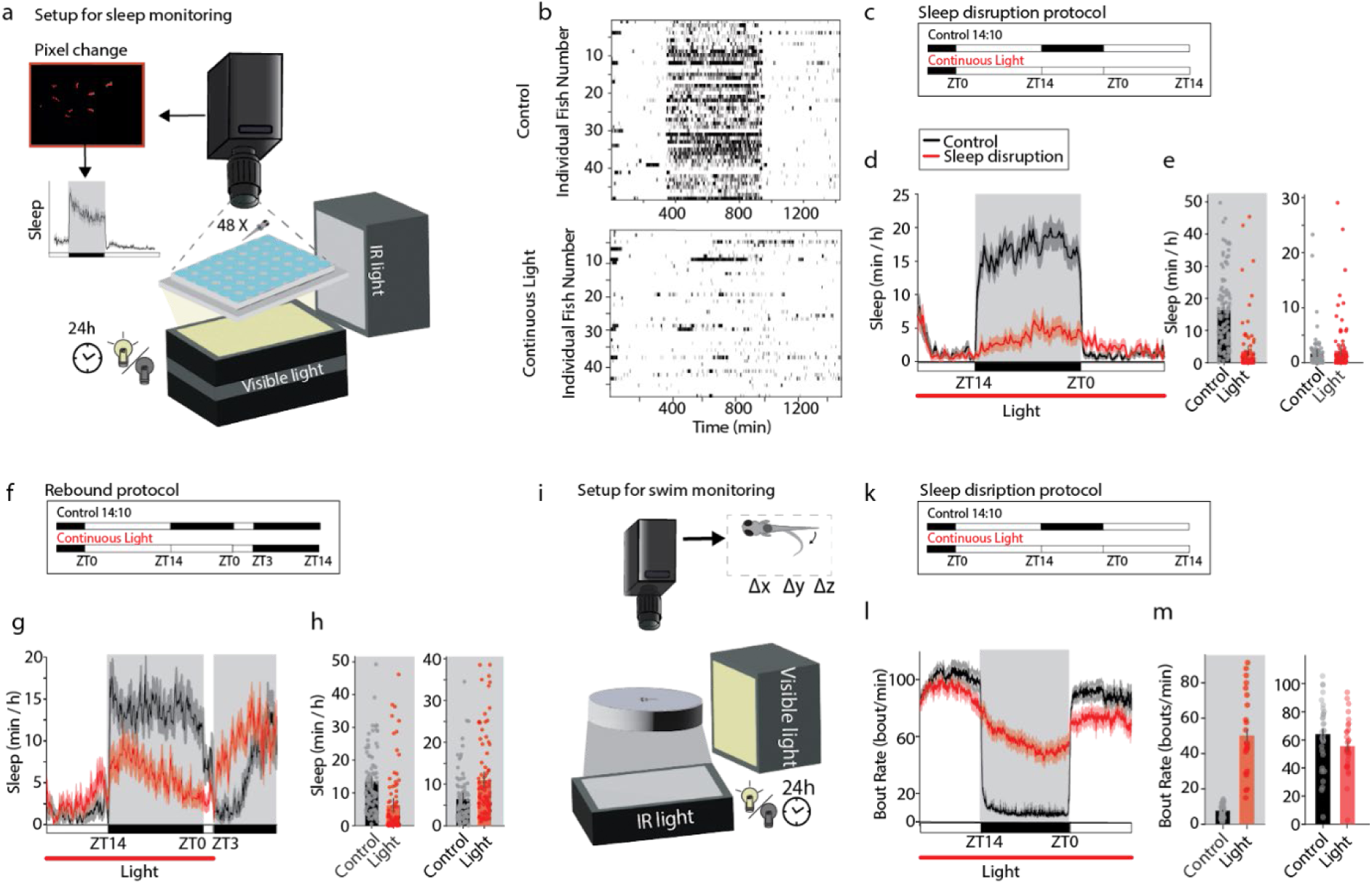
Sleep disruption in larval zebrafish using light. **a:** Schematics of the setup for sleep monitoring. Sleep of the individual was observed continuously for 24 h by determining pixel changes using an infrared camera. Visible light was introduced during the day and sleep disruption experiments. **b:** Representative sleep traces of individual fish during sleep disruption and control conditions. A black bar indicates that the animal was asleep for at least 1 min. n = 48/group. **c:** Sleep disruption protocol using continuous light. **d:** Lighting at night reduced the larvae’s sleep. **e:** In sleep-disturbed fish, sleep was decreased during the night (independent samples t-test: p < 0.001, t = 8.82, df = 168.68, n = 96/group) and slightly elevated on the day post-disruption (independent samples t-test: p = 0.34, t = −0.95, df = 196.47, n = 96/group). **f:** Larvae were exposed to complete darkness during the day following the sleep disruption. **g**: During daytime darkness, sleep-disrupted fish showed sleep rebound. **h:** During the sleep disruption, sleep was reduced compared to the control (independent samples t-test: p < 0.001, t = 5.08, df = 178.37, n_ctrl_ = 96, n_SD_= 96). Sleep time increased in the subsequent daytime darkness in disrupted fish (independent samples t-test: p < 0.001, t = −3.72, df = 150.83, n = 96/group). **i:** Schematics of the setup for swim monitoring. Bouting activity of the individual was observed continuously for 24 h using an infrared camera. **k:** Visible light was introduced during the day and sleep disruption experiments. **l:** Sleep-disrupted larvae increased their bout rate compared to controls. On the day post-disruption, their bout rate decreased. **m:** Constant light for 24 h increased the fish’s bout rate during nighttime (independent samples t-test: p < 0.001, t = −12.41, df = 31.53, n = 32/group) and decreased the bout rate during the day following sleep disruption (independent samples t-test: p = 0.008, t = 2.76, df = 61.58, n=32/group). Graphs show mean ± SEM with individual fish overlaid in circles.

Hypothesizing that daytime light may partially mask the sleep rebound ^26,38^, we assessed post-disruption sleep in darkness (Fig. 1f). As expected, sleep-disrupted fish showed a more pronounced rebound during a subsequent daytime darkness period (Fig. 1g, h, Fig. S1e-h). These findings demonstrate that light exposure effectively disrupts sleep in larval zebrafish and elicits compensatory rebound sleep.

In addition to sleep tracking, we monitored locomotor activity over a 24 h period using a high-resolution camera that enabled precise tracking of swim bouts (Fig. 1i, k). Sleep-disrupted larvae displayed higher bout rates during nighttime light exposure and lower bout rates the following morning, consistent with our sleep quantifications (Fig. 1l, m, S1i-l).

### Sleep disruption improves correctness in visual decision-making task

To investigate how lack of sleep affects vision-based decision-making in larval zebrafish, we examined their innate optomotor response (OMR) following sleep disruption. We presented larvae with moving dot stimuli known to elicit OMR ^27^. Larvae were placed in a dish, with a dot-motion kinematogram projected onto the bottom (Fig. 2a). This visual stimulus was presented in a closed-loop manner, moving perpendicular to the fish’s head direction, with coherence levels of 0%, 25%, 50%, or 100% in random order. These coherence levels introduced varying levels of difficulty for the larvae: at 0% coherence, all dots moved randomly, allowing for spontaneous swimming, while at 100% coherence, all dots moved uniformly, triggering the strongest OMR response. Intermediate coherence levels (25% and 50%) had subsets of dots moving in the same direction while the rest moved randomly. Using a high-resolution camera to track movements, we quantified bout rate, turning angles, reaction time, and performance, calculated as the likelihood of turning in the direction of the whole-field motion to stabilize their position in space (Fig. 2b). As expected, performance improved with increasing coherence due to greater directionality in the visual stimulus. Surprisingly, sleep disruption improved performance across all coherence levels (Fig. 2b, Fig. S2a, b) and reduced errors compared to rested sibling controls (Fig. S2c). This improved performance was observed with continuous, pulsed, and 6 h light exposures (Fig. S2a-l). Notably, these improved performances could not be attributed to inherent differences, as the probability for correct turns was comparable across all sibling fish on the day before sleep disruption (Fig. S2m-p).

**Figure 2:**
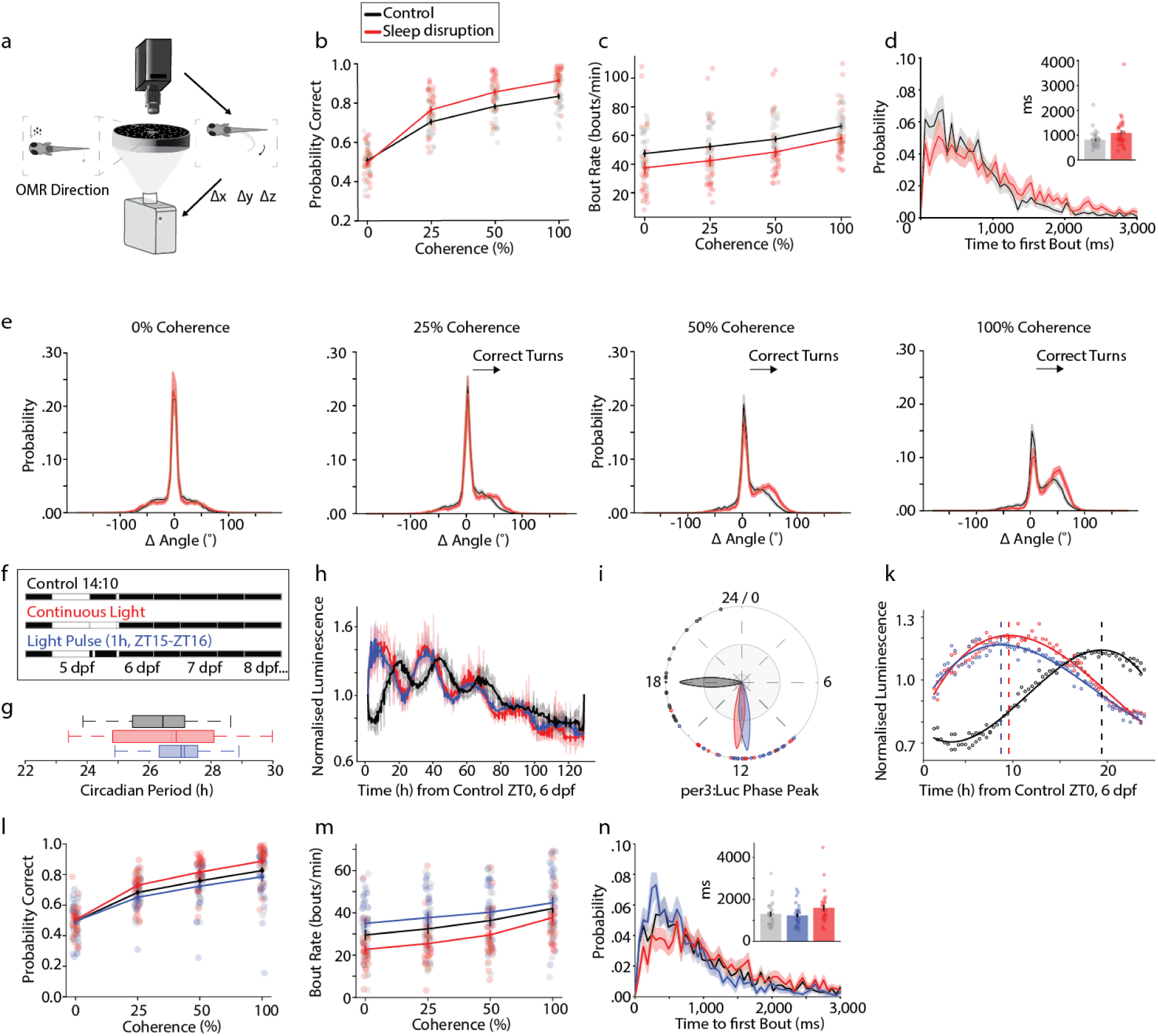
Sleep disruption but not a circadian phase shift increased performance and reaction time in visual decision-making task. **a:** The optomotor response of individual fish was observed by projecting a dot-motion kinematogram onto the bottom of the dish. The kinematogram consisted of dots moving with 0%, 25%, 50%, or 100% coherence perpendicular to the fish. A camera allowed tracking of fish in real time ensuring stimulus presentation in a closed-loop fashion. **b:** Performance, measured as the probability to make a correct turn in the direction of stimulus movement, was increased in sleep-disrupted fish compared to non-disrupted fish (mixed ANOVA: p < 0.001, F (1,61) = 12.94; Tukey’s HSD: p_0%_ = 0.39, p_25%_ = 0.004, p_50%_ < 0.001, p_100%_ < 0.001; n = 32/group). **c:** Sleep-disrupted fish showed a significantly reduced bout rate during the optomotor task (mixed ANOVA: p = 0.03, F(1,61) = 4.8; Tukey’s HSD: p_0%_ = 0.048, p_25%_ = 0.038, p_50%_ = 0.036, p_100%_ = 0.024, n=32/group). **d:** The reaction time after the onset of the 100% coherent stimulus was increased in disrupted fish. Inset shows the mean time to the first bout (independent samples t-test: p = 0.038, t = −2.13, df = 61, n = 32/group). **e:** Correct turns are displayed as turns with a positive Δ angle to the larvae’s previous position while incorrect turns are displayed as a negative Δ angle. Straight swims with a turning angle of −2.5° to 2.5° were most probable for all stimuli strengths and occurred independently of disruption. Turning angle distribution was similar between groups for spontaneous swimming (0% coherence) but not for directional stimuli. Disrupted fish made larger angle turns with increasing correctness. **f-k:** Circadian phase shift experiments. **f:** A 1 h light pulse from ZT15-16 was used to mimic the circadian effects of constant light at night as shown by per3:Luc bioluminescence measurements. **g:** Circadian period (solid line = mean, dashed line = median, box = interquartile range, whiskers = max/min) is not affected. **h:** per3:Luc bioluminescence across 120 h in constant dark after light treatments. **i:** Circadian per3:Luc phase peak, plotted as divisions of the circadian period, during hours 24-130 after ZT0 following light treatments (g-i, n_ctrl_ = 19, n_cont_ =16, n_1h-lp_ = 20). **k:** per3:Luc bioluminescence during first 24 h after ZT0 following light treatments, when behavioral decision-making tasks were performed. Phase peak indicated by dashed vertical line. Both the constant light and 1h light pulse conditions show a similar phase advance relative to the control group (control - constant light = 9.8 h, control - light pulse = 10.6 h. n_ctrl_ = 87, n_cont_ = 82, n_1h-lp_ = 89). **l-n:** Larvae were tested in the OMR paradigm either after a regular dark night, a single light stimulus from 12 AM to 1 AM or constant light for one night. **l:** The probability to make a correct turn improved in fish sleep deprived with constant light (mixed ANOVA: p = 0.015, F (1,53) = 6.26; Tukey’s HSD: p_0%_ = 0.43, p_25%_ = 0.016, p_50%_ = 0.052, p_100%_ = 0.0045; n = 28/group) but not in fish exposed to a 1 h light stimulus (mixed ANOVA: p = 0.0205, F (1,53) = 1.64, n= 28/group). **m:** The bout rate decreased in fish subjected to constant light in the previous night compared to the control (mixed ANOVA: p = 0.059, F (1,53) = 3.71, n= 28/group), but increased slightly in fish subjected to a 1 h light stimulus (mixed ANOVA: p = 0.207, F (1,53) = 1.62, n= 28/group). **n:** For a 100% coherent stimulus, sleep-disrupted fish showed a longer reaction time, (independent samples t-test: p = 0.14, t = −1.50, df = 53, n = 28/group), while fish subjected to a 1 h light stimulus did not show a change in reaction time (independent samples t-test: p = 0.62, t = 0.49, df = 53, n = 28/group). Graphs show mean ± SEM with individual fish overlaid in circles.

To investigate how this enhanced performance arises, we quantified larval swimming properties. Since bout rates were lower the morning following sleep disruption (Fig. 1l, m), we hypothesized that larvae would also exhibit reduced bout rates during the OMR task. As expected, sleep-disrupted animals consistently showed lower bout rates than controls across all stimulus coherences (Fig. 2c, Fig. S2e, i), alongside longer reaction times after stimulus onset (time_ctrl_ = 813 ± 60 ms [mean ± SEM], time_SD_ = 1083 ± 113 ms [mean ± SEM]) (Fig. 2d, Fig. S2f, k). Additionally, we observed larger turning angles in sleep-disrupted fish in response to coherent stimuli compared to the control group (Fig. 2e, Fig. S2g, l).

These findings align with current visual circuit models ^27,33^ which suggest that visual stimulus integration over time supports OMR. In our previous analyses of OMR behavior we have found improved performance correlated with longer response times (time to first bout after stimulus onset) ^38^, indicating that sleep disruption may affect decision-making through altered bout rate and reaction time. Supporting this speed-accuracy trade-off notion, our correlation analysis (Fig. S3a) (Spearman’s rank correlation coefficient, see Methods) revealed a negative correlation between bout rate and correct turning decisions (Spearman’s rank correlation coefficient: *control*: ρ = - 0.43, p = 3.9 e^-7^; *SD:* ρ = - 0.36, p = 4.1 e^-5^, n = 48/group).

Despite the increased accuracy in discriminating visual stimuli, sleep disruption and the resulting extended decision time likely have adverse effects on larvae. For instance, the reduced bout rate in the OMR task suggests decreased exploratory behavior, with sleep-disrupted fish covering less distance than their non-disrupted siblings (Fig. S3b-f).

Together, these results suggest that light-induced sleep disruption slows down the larvae’s reaction time, allowing larvae to integrate the visual stimulus for longer and thereby increase correct turns.

Since abnormal light exposure at night is known to affect the phase timing of the endogenous circadian clock, we tested whether altered circadian rhythms could contribute to improved OMR performance. Using a luciferase reporter of circadian gene expression (per3:Luc, see Methods), we observed that both constant light and a 1 h light pulse delivered 1 h after night onset (ZT15) advanced the circadian rhythm by approximately 10 h compared to the control group, which remained in darkness throughout the night (Fig. 2f-k). Although both lighting conditions induced a similar phase shift, a 1 h light pulse caused minimal sleep disruption (Fig. 2h). Unlike constant light exposure, the 1 h light pulse did not enhance OMR performance compared to controls (Fig. 2l), nor did it reduce bout rate (Fig. 2m) or lengthen reaction time [time_ctrl_ = 1300 ± 117 ms, time_SD_ = 1588 ± 150 ms, time_1h pulse_ = 1224 ± 99 ms] (Fig. 2n).

These findings suggest that circadian phase shifts induced by light do not alone account for improved OMR performance, but rather that these enhancements are due to other light-induced effects, for example reduced sleep time.

### Artificially prolonged reaction time improves optomotor performance

To further test how extended reaction times influence OMR performance, we administered melatonin to artificially increase reaction time. Previous studies have found that melatonin, a natural hormone that promotes sleep, dose-dependently reduces locomotion in zebrafish ^9,39,40^. Therefore, we hypothesized that melatonin would slow OMR reaction times and consequently enhance task correctness.

As predicted, melatonin dose-dependently enhanced OMR performance, with higher concentrations decreasing bout rate and improving correctness relative to pre-treatment and vehicle controls (Fig. 3a, S4a-l). This effect was reversible upon melatonin washout (Fig. S4b). Specifically, 100 nM melatonin improved OMR performance (Fig. 3b) and reduced error rate (Fig. S4c). Likewise, following melatonin treatment, bout rate decreased (Fig. 3c), and reaction time increased (time_ctrl_ = 922 ± 156 ms [mean ± SEM], time_mel_ = 1699 ± 143 ms [mean ± SEM]) (Fig. 3d) supporting a speed-accuracy trade-off hypothesis. Additionally, melatonin-treated fish displayed larger angle turns toward coherent stimuli while straight swims were unaffected, similar to sleep-disrupted fish (Fig. 3e).

**Figure 3:**
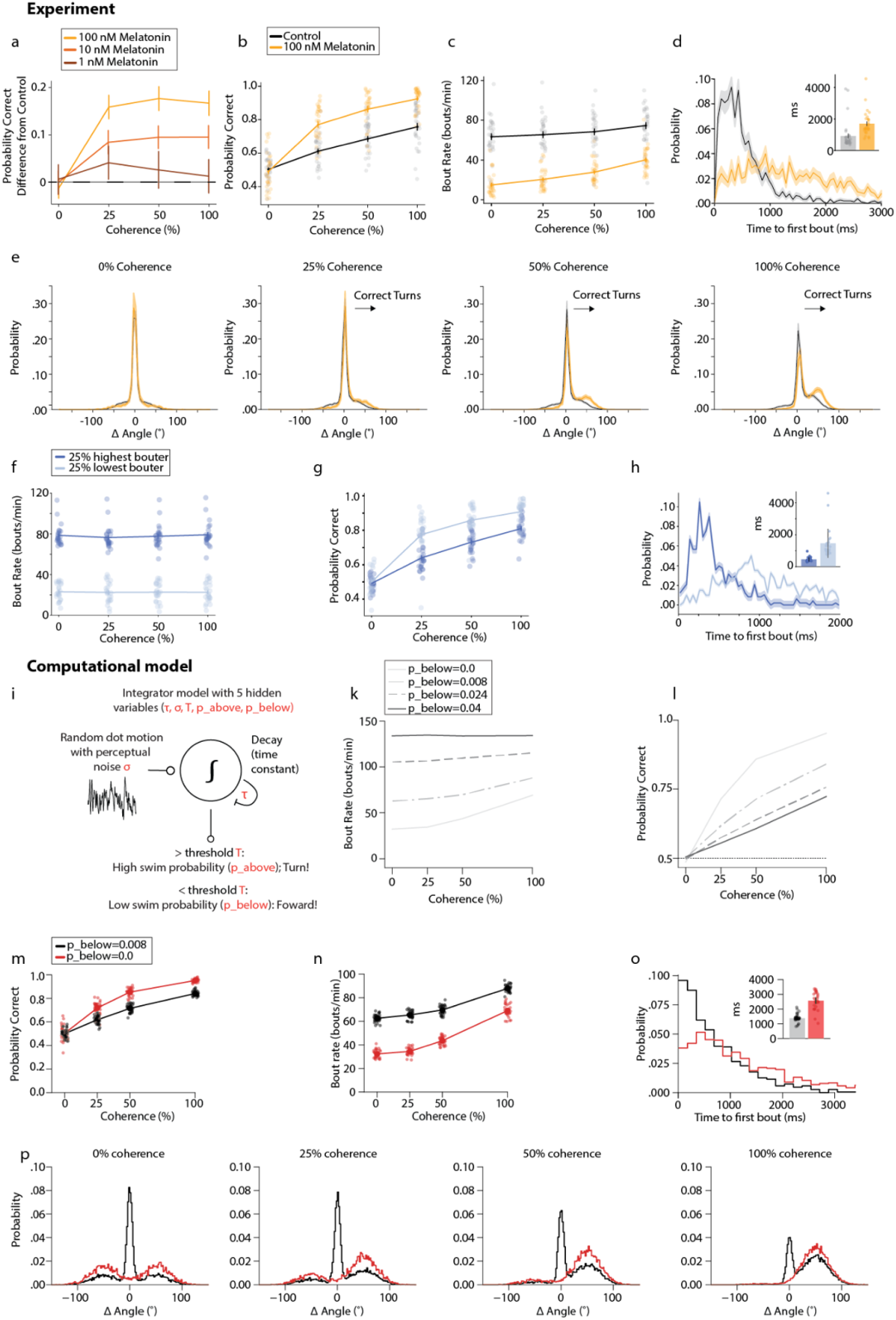
Longer reaction time leads to increased OMR performance in decision-making task. **a:** Melatonin treatments led to better OMR performances in a dose-dependent manner compared to control trials in the same larvae. n_100 nM_ = 32/group, n_10 nM_ = 32/group, n_1 nM_ = 16/group. **b:** The fish’s probability of correct turns increased after administration of 100 nM melatonin compared to trials before melatonin treatment (repeated measures ANOVA: p < 0.001, F (1,29) = 52.68; Tukey’s HSD: p_0%_ = 0.27, p_25%_ <0.001, p_50%_ < 0.001, p_100%_ < 0.001, n = 32). **c:** The bout rate was decreased after treatment with 100 nM melatonin compared to before treatment (repeated measures ANOVA: p < 0.001, F (1,29) = 111.33; Tukey’s HSD: p_0%_ < 0.001, p_25%_ <0.001, p_50%_ < 0.001, p_100%_ < 0.001, n = 32). **d:** Reaction time after onset of the 100% coherent stimulus was slower after melatonin treatment. Inset shows the mean time to the first bout (paired samples t-test: p = 0.004, t = −3.13, df = 29, n = 32). **e:** The turning angle distribution was similar before and after melatonin treatment for spontaneous swimming (0% coherence), but melatonin increased larger angle turns and improved the fish’s performance for directional stimuli. **f:** Non-perturbed siblings with innately low bout rates and innately high bout rates were seperated (mixed ANOVA: p < 0.001, F (1,46) = 362.59; Tukey’s HSD: p_0%_ < 0.001, p_25%_ < 0.001, p_50%_ < 0.001, p_100%_ < 0.001; n = 24/group). **g:** Individuals with an innately low bout rate showed improved OMR performance compared to innately highly bouting fish (mixed ANOVA: p < 0.001, F (1,46) = 31.92; Tukey’s HSD: p_0%_ = 0.63, p_25%_ < 0.001, p_50%_ < 0.001, p_100%_ < 0.001; n = 24/group). **h:** The fish with the innately lowest bout rates had slower reaction times compared to the fish with the innately highest bout rates (independent samples t-test: p < 0.001, t = −5.22, df = 46, n = 24/group). **i:** Integrator model. Figure adapted from Harpaz et al. (2021) ^33^. **k:** Decreasing bouting probability for forward swims (p_below_) caused decreased bout rates and improved performances (**l**), reflecting the observed behavioral phenotype. **m-p:** Computationally decreasing p_below_ from 0.008 (black) to 0 (red) mimicked the behavioral phenotype after sleep disruption. n_ctrl_=32, n_SD_ = 32. **m:** Decreasing p_below_ to 0, increased the agent’s probability for correct turns. **n:** Decreasing p_below_ reduced the bout rate. **o:** The reaction time was slowed down with a decreased probability for straight swims. **p:** Probability distribution of turn angles for 4 different coherence levels. Note that for p_below_ = 0, the probability of unbiased forward swims is zero, and therefore a peak around 0 degree was not observed. Graphs show mean ± SEM with individual fish/agents overlaid in circles.

If bout rate predicts performance, we would expect to see a correlation between bout rate and correctness in untreated fish (Fig. 3f-h). Thus, innately low bouting fish should exhibit increased performance compared to high bouting fish. When we compared the 25% of highest-bouting non-perturbed larvae to the 25% of the lowest-bouting non-perturbed larvae (Fig. 3f), we indeed observed that the lowest-bouting 25% made more correct turns than high-bouting counterparts (Fig. 3g). Correspondingly, we observed slower reaction times in the low-bouters compared to the high-bouters (time_high_ = 460 ± 144 ms [mean ± SEM], time_low_ = 1461 ± 908 ms [mean ± SEM]) (Fig. 3h). Likewise, low-bouters exhibited greater turning angles than high-bouters (Fig. S4m). Together, these results suggest that increased reaction time and reduced bout rate are linked to improved OMR performance, even without sleep disruption.

### Downregulation of motor command probabilities for spontaneous swims increases OMR performance

To pinpoint circuit elements potentially underlying improved performance, we employed computational modeling with a drift diffusion model for motion integration ^27,33^ (Fig. 3i). The bounded leaky integrator circuit model consists of an integrator (∫) that accumulates noisy (σ) coherence evidence (c) over time. The integrator is leaky (τ) and slowly decays to 0 without stimulation. If the integrated value falls below a threshold (T), a forward swim is initiated with probability p_below_; if it exceeds T, a turning motor command is relayed with probability p_above_. We hypothesized that sleep disruption could alter sensory-motor convergence by affecting one of five parameters:

1. Increased Threshold (T): Higher thresholds would prolong accumulation of the visual stimulus, resulting in more forward swims and fewer turns, ultimately reducing correct decisions. Modeling an elevated threshold decreased swim rates (Fig. S5b) and, as expected, reduced the probability of correct turns (Fig. S5c), a result incompatible with our data showing reduced bout rate and increased performance (Fig. 2b, d; Fig. 3a-c), ruling out threshold increases.
2. Extended Decay Constant (τ): Increasing τ would enable longer visual integration, filtering out noise more strongly and thereby enhancing performance. However, simulations revealed only a minor improvement in correct turns, with lower bout rates and slower response times (Fig. S5d, e), excluding this as a potential mechanism.
3. Noise Variability (σ): Changes in internal perceptual noise of the dot motion integration could affect turning probability. Although increased or decreased noise altered bout rate, it did not increase correct turns (Fig. S5f, g), ruling out noise as a mechanism for improved accuracy in sleep-deprived fish.
4. Lower Turn Probability (p_above_): Reduced p_above_ would decrease turn accuracy, contrary to the observed behavioral improvements, leading us to exclude this explanation (Fig. S5h, i).
5. Reduced Spontaneous Swim Probability (p_below_): Lowered p_below_ would decrease swim rates and enhance decision accuracy, matching our experimental data. This explanation accounts for both overnight sleep disruption with light (Fig. 2b, d) as well as melatonin-induced alterations of bouting (Fig. 3a-c). Accordingly, we propose that sleep disruption and melatonin-treatment modulate the motor command probabilities for spontaneous swims. Simulating decreased p_below_ from 0.008 (control) to 0 (sleep-disrupted) aligned with observed behavioral outcomes: increased accuracy (Fig. 3m), reduced bout rates (Fig. 3n), elongated reaction times (Fig. 3o), and increased turning angles (Fig. 3p).

In conclusion, the computational modulation of the swimming probabilities confirmed that slowed bouting increases the fish’s probability for correct turns through a slowed reaction time.

### Sleep disruption increases avoidance of aversive odors

To assess whether sleep loss affects sensory performance across multiple modalities, we tested sleep-disrupted fish on gradients of aversive odors in a known decision-making task ^34^ (Fig. 4a). Using each fish’s position, we calculated a preference index (PI) for cadaverine and putrescine, with positive PI values indicating attraction and negative PI values indicating aversion. As anticipated, control fish showed greater avoidance (negative PI) at higher cadaverine concentrations, spending more time away from stronger odor zones (Fig. 4b). After sleep disruption, avoidance of high cadaverine concentrations increased (Fig. 4c), and a similar trend was observed with putrescine, suggesting a general increase in aversion to unpleasant odors (Fig. 4d).

**Figure 4:**
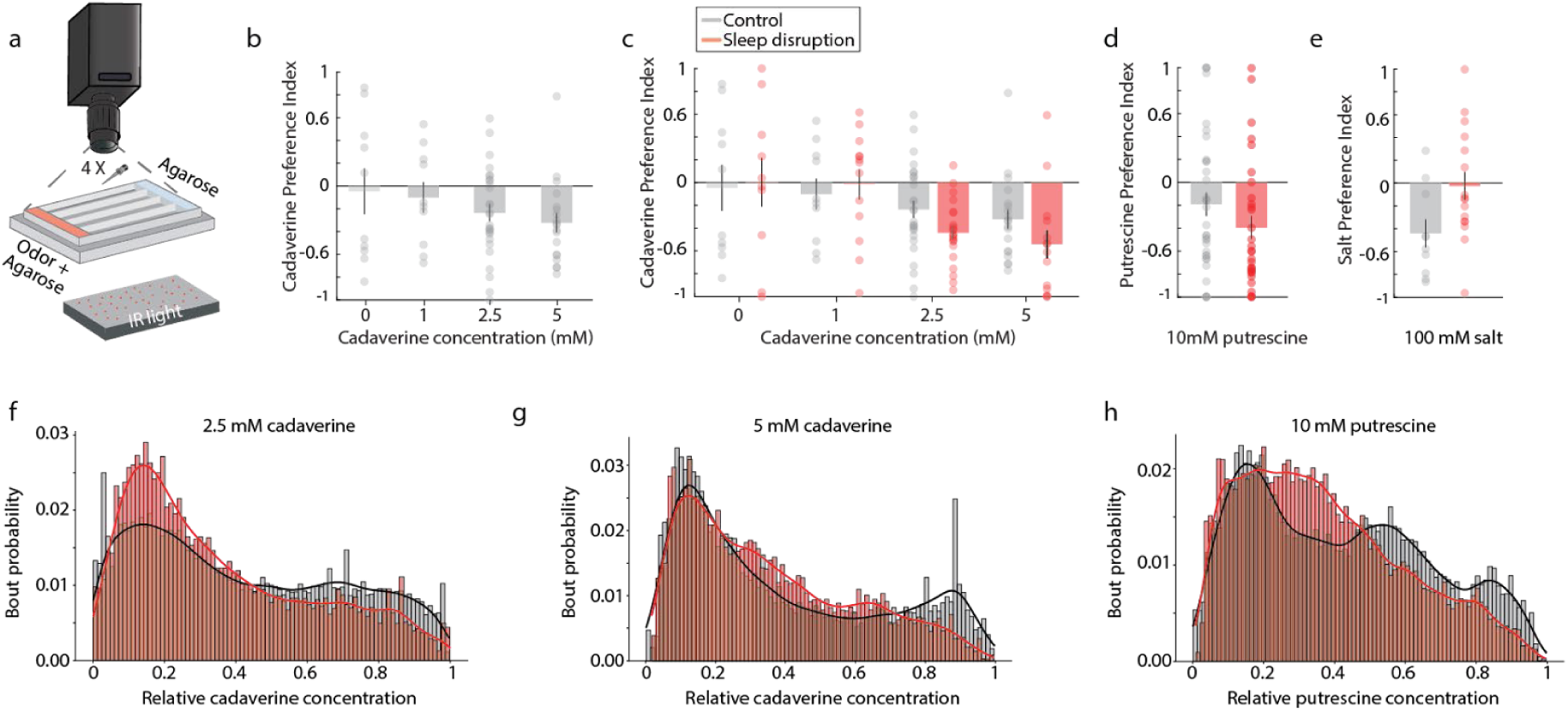
Sleep disruption increased avoidance of aversive odors. **a:** Scent gradients were created across 4 individual closed-off lanes of water. The position and swimming behavior of individual fish within each lane were observed under visible and IR light for 30 min using an infrared camera. **b:** A preference index (PI) was calculated for each fish summarizing position over the duration of each experiment. Fish avoided high cadaverine concentrations, resulting in negative PIs. n_0mM_ = 10, n_1mM_ = 10, n_2.5mM_ = 27, n_5mM_ = 19. **c:** Sleep-disrupted fish exhibited an increased avoidance of cadaverine compared to sibling controls (Wilcoxon rank sum test: p_0mM_ = 0.88, p_1mM_ = 0.64, p_2.5mM_ = 0.04, p_5mM_ =0.14; light: n_0mM_ = 10, n_1mM_ = 13, n_2.5mM_ = 22, n_5mM_ = 14). The control group in c is the same as in b. **d:** Sleep-disrupted fish also slightly increased their avoidance of putrescine (Wilcoxon rank sum test: p = 0.18, n_ctrl_ = 33, n_SD_ = 30). **e:** Sleep-disrupted fish show a reduced aversion to 100 mM salt (Wilcoxon rank sum test: p = 0.037, n_ctrl_ = 10, n_SD_ = 16). **f:** In 2.5 mM cadaverine, sleep-disrupted fish showed an earlier return point through a decreased bout count in high odor concentrations. **g:** In 5 mM cadaverine, sleep-disrupted fish returned to low cadaverine concentrations earlier than control fish. **h:** Sleep-disrupted fish spent less time in high putrescine concentrations than controls, characterized by a decrease in bout count close to the putrescine source. Bar graphs represent mean ± SEM with the responses of individual fish overlaid in circles. Histograms show the normalized bout count probabilities across 85 bins including a kernel density estimate.

To determine if heightened avoidance was specific to odors, we also tested responses to high salt concentrations. Unlike their reaction to odors, sleep-disrupted fish showed reduced avoidance of 100 mM NaCl, suggesting that sleep disruption selectively enhances aversion to odors rather than to all aversive stimuli (Fig. 4e).

We investigated potential behavioral strategies underlying this increased avoidance of aversive odors, considering three possibilities:

1. Fish could increase their tuning angle (turn more sharply) when encountering an aversive odor to facilitate escape. However, turning angle analyses showed that neither control nor sleep-disrupted fish altered their turning angles in response to high odor concentrations (Fig. S6a-d).
2. Fish could increase their bout rate near the odor source. Yet, bout rate analyses between the odor and control areas showed no significant increase in bout rate after sleep disruption, for both cadaverine and putrescine. Bout rates decreased in sleep-disrupted animals when fish were exposed to high salt concentrations and increased in well-rested siblings (Fig. S6e-g).
3. Fish could turn away from aversive odors earlier, reducing time spent in areas of high odor concentration. We assessed bout distribution along the cadaverine gradient and found that bouts were evenly distributed in odorless conditions (Fig. S6h). However, as cadaverine concentration increased, bout count decreased for both experimental groups but especially in sleep-disrupted fish, suggesting that they avoided high concentrations by turning away sooner (Fig. 4f, g, S6i). This behavior was more pronounced at higher cadaverine concentrations (2.5 mM and 5 mM), with sleep-disrupted fish showing a leftward shift in their return point along the gradient, indicating heightened odor sensitivity following sleep disruption. This effect was consistent for putrescine (Fig. 4h) but absent for salt, with sleep-disrupted fish showing an evenly distributed bout count in high and low salt areas, while control fish bout counts were shifted towards low salt concentrations (Fig. S6k).

In summary, these results indicate that sleep-disrupted larvae exhibit a stronger odor avoidance by shifting their turning point and thus turning back earlier when encountering aversive odors. This response is specific to odors and does not extend to other aversive stimuli like salt.

### Low-dose cortisol mimics sleep disruption in aversive odor avoidance

To investigate the circuit mechanisms behind increased odor avoidance following sleep disruption, we tested two hypotheses from the literature:

First, sleep disruption might induce acute starvation, which could affect olfactory sensitivity and thus increase avoidance behavior. Although studies in humans and mammals suggest contradictory effects, some have linked increased caloric intake ^41–43^ and energy expenditure to sleep deprivation ^44,45^. Yet, despite observing a slight change in bout rate in our experiments after complete starvation, we dismissed this hypothesis, as neither partial nor complete starvation mirrored the increased cadaverine avoidance or return point changes seen in sleep-disrupted fish (Fig. S7a-e).

A second explanation for increased avoidance is that sleep disruption induces stress which heightens sensitivity to aversive odors. Previous studies have shown that higher cortisol levels are associated with enhanced olfaction ^48,49^. To test whether stress from sleep loss improves odor-based decision-making, we measured internal cortisol levels in sleep-deprived fish using a luminescence assay (Fig. 5a). Due to variability in cortisol levels in control fish (Fig. S7f), we calculated the relative cortisol increase in sleep-disrupted fish by subtracting the cortisol levels of well-rested fish from their sleep-disrupted siblings. As expected, sleep-disrupted fish showed elevated cortisol levels, suggesting that sleep disruption causes stress (Fig. 5a).

**Figure 5:**
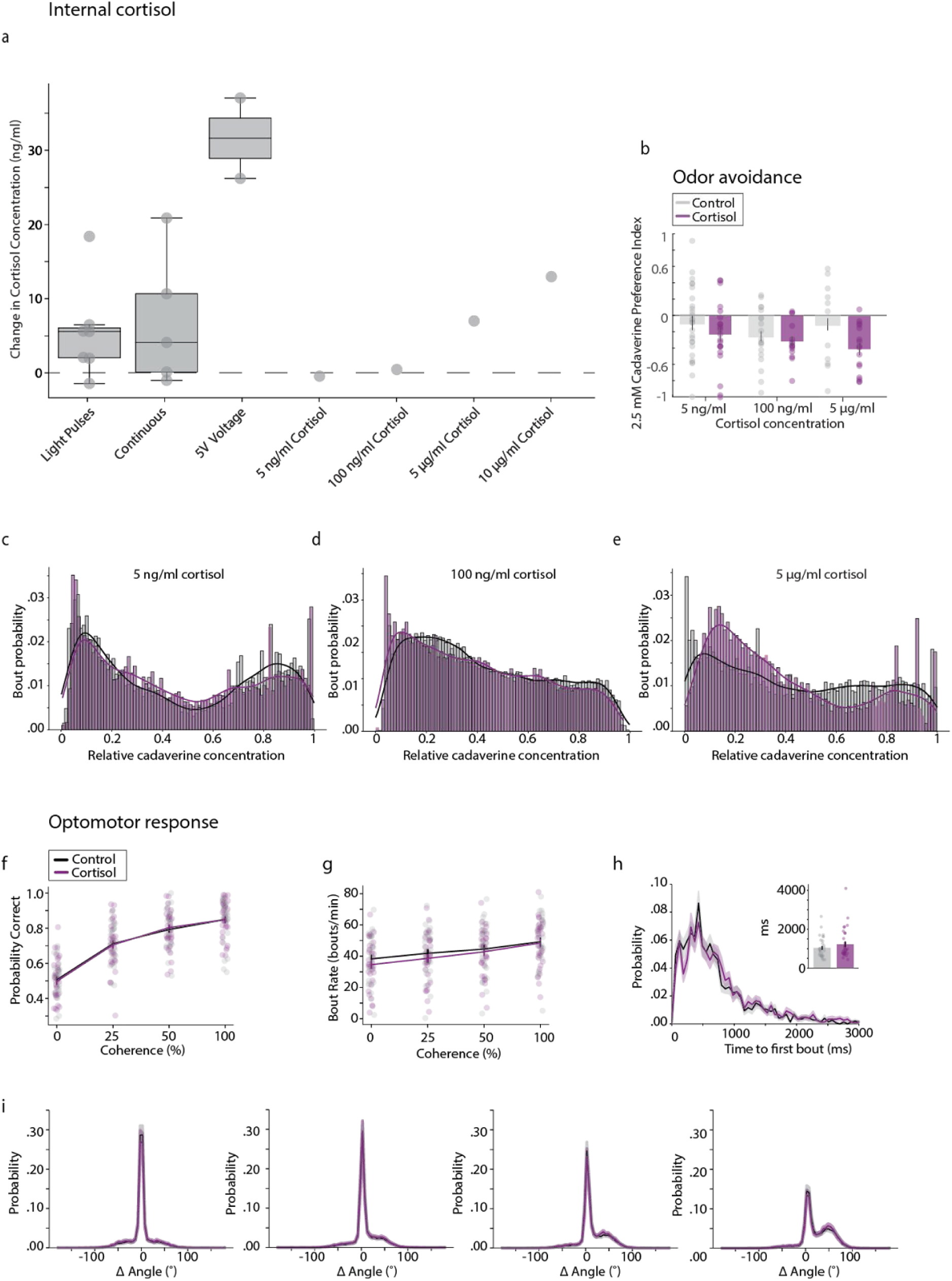
Cortisol treatment can mimic behavioral changes in olfactory-based, but not visual-based decision-making tasks. **a:** Internal cortisol levels were increased in sleep-disrupted fish compared to sibling controls (independent samples t-test: *light pulses*: p = 0.031, t = −2.06, df = 12, n = 7; *continuous*: p = 0.043, t = −1.9, df = 10, n = 5). An acute 5V shock increased cortisol levels more than sleep disruption (independent samples t-test: *5V voltage*: p = 0.02, t = −4.88, df = 2, n = 2). Treatment with external cortisol (5 ng/ml, 100 ng/ml, 5 µg/ml, and 10 µg/ml) for 30 minutes increased internal cortisol levels in a concentration dependent manner (n = 1). Each circle represents a group of 30 larvae. **b-e:** The effects of cortisol treatment on larval zebrafish behavior in the olfactory decision-making task. **b:** Treatment with 5 µg/ml of cortisol elicited more negative PIs for 2.5 mM cadaverine compared to control. Treatment with lower doses of cortisol (5 ng/ml and 100 ng/ml) showed no discernible changes in behavior (Wilcoxon rank sum test: *5 ng/ml cortisol:* p = 0.26, n_ctrl_ = 27, n_cort_ = 21; *100 ng/ml cortisol*: p = 0.46, n_ctrll_ =19, n_cort_ = 15; *5 µg/ml cortisol*: p = 0.06, n_ctrl_ = 11, n_cort_ = 16). **c:** Fish treated with 5 ng/ml cortisol, showed a marginally reduced bout count in high cadaverine concentrations. **d:** Treatment with 100 ng/ml cortisol did not alter the larvae’s bout probability distribution along the cadvaerine gradient. **e:** Treatment with 5 µg/ml cortisol reduced the bout probability in high odor concentrations, indicating an earlier return point. **f - i:** The effects of treatment with 5 µg/ml cortisol on larval zebrafish behavior in the visual decision-making task. **f:** Cortisol treatment did not improve the fish’s performance in the optomotor task (repeated measures ANOVA: p = 0.89, F (1,61) = 0.02, n= 28/group). **g:** Cortisol treatment had no effect on the bout rates (repeated measures ANOVA: p = 0.56, F (1,62) = 0.35, n = 28/group). **h:** Cortisol treatment did not elicit changes in reaction time (independent samples t-test: p = 0.24, t = −1.17, df = 62, n=28/group). **i:** Turning angle distributions remained unchanged after cortisol treatment. Bar graphs represent mean ± SEM with the responses of individual fish overlaid in circles. Histograms show the normalized bout count probabilities across 85 bins including a kernel density estimate.

To contextualize the stress response after sleep disruption, we included a group subjected to an acute stressor (a 5V electric shock) ^50^, which raised cortisol levels 3-5 times higher than those observed with sleep disruption, suggesting that sleep loss induces mild stress (Fig. 5a).

Next, we investigated whether this stress response underlies increased aversive odor avoidance. To test this, we activated the fish’s stress response by adding water-soluble cortisol ^50^ at various concentrations (5 ng/ml, 100 ng/ml and 5 µg/ml) (Fig. 5a) to match the internal cortisol levels seen after sleep disruption. When measuring the internal cortisol levels in these well-rested cortisol-treated fish, we discovered that exposure to 5 µg/ml of cortisol produced comparable cortisol levels as seen after sleep loss.

Subsequently, we examined how fish reacted to an aversive odor after external cortisol treatment. After cortisol treatment, fish showed stronger cadaverine avoidance, reflected in more negative PI values, with 5 µg/ml cortisol resulting in stronger aversion than untreated controls (Fig. 5b). Similarly, these cortisol-treated fish turned away from cadaverine earlier, avoiding high concentrations and decreasing their bout probability in areas of high odor concentration (Fig. 5c-e), closely mirroring sleep-disrupted behavior. This cortisol-induced behavior, like sleep disruption, did not affect heading angle (Fig. S6d) or bout rate (Fig. S7g), suggesting a shared avoidance strategy.

Finally, we examined whether cortisol also contributes to increased correctness after sleep disruption by testing OMR performance after treatment with 5 µg/ml cortisol. Unlike in the olfactory task, cortisol treatment did not alter performance in the OMR paradigm. Fish maintained the same probability to turn in the direction of dot motion (Fig. 5f) and a similar number of incorrect turns (Fig. S7h), and showed no change in bout rate (Fig. 5g) or reaction time (time_ctrl_ = 1039 ± 91 ms [mean ± SEM], time_cort_ = 1231 ± 134 ms [mean ± SEM]) (Fig. 5h). Likewise, no changes in the turning angle distributions were found post-treatment (Fig. 5i).

Together, these findings indicate that sleep disruption in larval zebrafish acts as a mild stressor, resulting in a heightened avoidance of aversive odors, likely due to increased sensitivity. Cortisol treatment alone was enough to replicate these effects in the olfactory task but not in the OMR, suggesting distinct underlying mechanisms for visual and olfactory task improvements after sleep disruption. In the visual OMR task, sleep disruption reduces bout rate, slowing reaction time and thus leading to more time for stimulus integration, while in the olfactory task, sleep disruption activates the stress response, enhancing odor aversion likely through increased odor sensitivity.

## DISCUSSION

We investigated how light-based sleep disruption affects decision-making in larval zebrafish. Our findings show that after sleep disruption, larval zebrafish improved their performance in two independent perceptual decision-making tasks, each improvement arising from independent mechanisms. In the OMR task, performance enhancement is attributed to a decreased spontaneous swim-bout rate and longer reaction times—an example of a speed-accuracy trade-off that permits extended stimulus integration time, resulting in more correct decisions. In the second task, larval zebrafish demonstrated increased odor avoidance, largely due to an earlier return upon encountering the aversive odor. Unlike in the OMR task, this heightened odor avoidance likely results from elevated cortisol levels post-sleep disruption. These results show that sleep disruption enhances performance via distinct, task-specific mechanisms. In the following, we discuss the implications and discuss these findings within the existing literature for each paradigm in sequence.

We were able to replicate the speed-accuracy trade-off effect observed after sleep disruption by treating zebrafish larvae with external melatonin, which is known to reduce swim bouting ^9,39,40^. Even though melatonin is likely depleted through light-induced sleep disruption ^51^, melatonin administration elicits similar behavioral effects in the OMR paradigm through its sleep-promoting effects. We observed performance improvement after melatonin treatment which supports the idea that the increased number of correct turns is caused by a slowing of locomotion speed and the resulting increase in stimulus integration time. Thus, we hypothesize that the speed-accuracy trade-off plays a pivotal role in enhancing OMR performance.

The relationship between sleep, stress ^24^ and melatonin levels is complex, with light timing, duration and intensity playing a role ^52–54^. However, our results show that light exposure at night strongly decreases sleep in larval zebrafish and elicits homeostatic sleep rebound the following day, consistent with other work in larval ^8,24^ but not adult ^22^ zebrafish. As expected ^26,51^, light exposure at night elicited a shift in the phase of the circadian clock (Fig. 2f-k). Importantly, while nighttime light exposure caused a shift in the circadian clock, this alone could not explain the improved OMR performance following sleep disruption by nighttime light (Fig. 2a-d), as shorter light pulses that induce equally strong phase advances in the circadian clock but cause relatively a small disruption of sleep at night failed to improve performance the next day (Fig. 2l-n). However, we cannot rule out other direct effects of light on zebrafish physiology ^55^ playing a role in the improved performance on visual and odor tasks.

In support of the argument that the increased OMR performance can be attributed to an increased integration time, computational modeling utilizing leaky integrator models ^27,33^ indicated that sleep disruption affects motor commands, resulting in prolonged reaction times, which in turn enhances performance (Fig. 3). Specifically, we identified a decrease in the spontaneous swimming probabilities as the most likely element of the drift diffusion model modulated by sleep disturbance. Along this line, in a separate study, we attributed increased OMR performance after sleep disruption to longer integration times ^38^, termed higher competence.

While we observed performance improvement after light-induced sleep disruption in the perceptual decision-making tasks, deterioration of performance as a consequence of sleep disturbance has been reported in many different species. Rodents ^52–54,56^, flies ^57,58^, and zebrafish ^51,59^ perform worse in a given task following sleep deprivation. In humans, sleep deprivation has been shown to deteriorate sensorimotor coupling ^60^. The effects of sleep deprivation on decision-making tasks, however, are mostly explored in human subjects which show impaired and more risky decision-making after extended wakefulness ^2,3,5^. Given that sleep disruption usually impairs decision-making, the performance improvements reported in this study were initially surprising. However, the notion of speed-accuracy trade-off in decision-making tasks has been reported across many species. Associations between slower but more accurate decisions have been found in zebrafish ^33,61^, insects ^62–66^, birds ^67^, mice ^68^, and humans ^69^ in visual as well as olfactory decision-making paradigms.

Here we report that in addition to the visual decision-making task, in an olfactory-based decision-making task, fish displayed increased avoidance of aversive odors following sleep disruption, which is largely explained by an earlier return upon encounter of the aversive odor. In contrast to the OMR task, the heightened odor avoidance likely stems from the elevated cortisol levels induced by sleep disruption, which demonstrates that sleep disruption can improve performance via multiple mechanisms.

Despite the difference in neuronal integration between vision and olfaction ^70^, we showed that performance in the olfactory avoidance task was enhanced as well after sleep disruption. This is opposing the literature, which commonly associates olfactory impairment with lack of sleep ^71–73^. However, it agrees with findings of increased reactivity to negative stimuli after sleep deprivation. Humans showed increases in amygdala activity to negative visual emotional stimuli ^74,75^ and falsely interpreted non-threatening stimuli as threatening ^76^.

In contrast to the vision-based decision-making task, where sleep-disrupted fish exhibited a reduced swim rate and slowed responses, in the odor-based decision-making task, fish showed no unambiguous change in bout rates in high-odor concentrations (Fig. S6e-g). Unlike in visual tasks, several experimental studies across different species suggest that olfaction does not profit from longer integration times ^77,78^. Therefore, it is likely that sleep disruption affects the two different sensory modalities of vision and olfaction in different manners. Whether sleep disruption heightens the fish’s odor sensitivity or simply their reaction to the odor remains to be investigated. Regardless, it appears that the earlier return in sleep-deprived fish supports the theory of an increased olfactory sensitivity following the lack of sleep. Moreover, both sleep-deprived and well-rested siblings increased their bout rate in the presence of the aversive odor (Fig. S6), possibly to facilitate a quicker escape from the odor signaling danger. This finding is in agreement with a previous study that discovered increased bout rate and faster turns to avoid a cadaverine stimulus ^36^.

In our study, we indicate that the increased olfactory sensitivity after sleep disruption is caused by increased cortisol levels. Increase in cortisol could function as a mechanism to heighten avoidance of potentially dangerous odors in an already stressful environment. In agreement with that, we found increased cortisol levels after sleep disruption in the larval zebrafish. Additionally, we could replicate the enhanced odor avoidance we observed after sleep disruption (Fig. 4) by external administration of cortisol (Fig. 5b-e). Therefore, we postulate that increased olfactory sensitivity is cortisol-mediated. Further testing using cortisol synthesis blockers like ketoconazole or metyrapone could improve our understanding of the role cortisol plays in modulating olfactory avoidance behavior.

While in adult zebrafish no changes in cortisol levels have been reported after extended light exposure ^79^, the effects of sleep disruption on cortisol levels in larval zebrafish had not been studied before. In humans, contradictory results have been reported regarding the effects of sleep disruption on cortisol levels ^81–84^. However, increased cortisol levels have been linked to improved olfaction in humans ^48,85^. Specifically, increased saliva cortisol levels were correlated with increased odor identification and decreased odor detection thresholds.

Additionally, our findings support the notion that sleep disruption using light exposure acts as a mild stressor compared to other sleep disruption methods like electrical shocks. Internal cortisol levels show moderate increases following light exposure while a single electrical shock at an intensity similar to the ones used in previous adult zebrafish sleep deprivation studies ^22^ elicits a significantly stronger cortisol elevation (Fig. 5a). However, further studies comparing internal cortisol levels in fish sleep deprived with electrical shocks or with mechanical shaking would be necessary to consolidate our hypothesis of light exposure as a mild stressor.

In the future, our research could help to systematically analyze how sleep disruption affects cortisol levels across vertebrates and help uncover the mechanisms behind enhanced olfactory sensitivity following sleep deprivation. We hypothesize that cortisol treatment in fish may further influence cognitive functions like attention.

Together, our findings reveal the surprising effects of sleep disruption on perceptual decision-making in larval zebrafish. Both visual- and olfactory-based decision-making improve following sleep disruption, yet through distinct mechanisms: visual decision-making benefits from a speed-accuracy trade-off driven by extended reaction times, while olfactory decision-making likely gains from cortisol-mediated stress responses. This study highlights how sleep disruption uniquely impacts different sensory modalities, with potentially profound implications for brain function. These findings lay important groundwork for future research into the brain circuits underlying sleep disruption’s effects.

## METHODS

### Zebrafish Husbandry

Adult fish were housed in the Harvard Biolabs fish facility and were crossed in pairs to obtain sibling controls. Eggs were collected 0 days post fertilization and raised in groups of ∼ 30 zebrafish larvae under a 14 h light/10 h dark cycle in a petri dish (15 cm in diameter) in filtered fish facility water at 28 °C. Every clutch was fed once a day with paramecia from 5 days post fertilization (dpf). Half of the water was exchanged daily and dead fish and chorions were removed. For all experiments, except Fig. S1b-h, larvae of the AB strain were used and entered the experiments between 5 - 8 days post fertilization (dpf). At this age, the sex cannot be determined. In Fig. S1b-h, WIK (6-7 dpf) were used. All experiments were conducted according to the guidelines of the Harvard Institutional Animal Care and Use Committee.

### Sleep Disruption

#### Sleep Monitoring

Sleep disruption was performed using light. To determine sleep the fish’s locomotor activity was tracked using a setup described in Joo et al. (2021) ^86^. In short, fish were tracked using a camera (Grasshopper3, NIR-FLIR system) with a 50 mm fixed focal lens with an IR filter and an IR light mounted beneath. A custom LabView program allowed precise timing of lighting conditions for the duration of the experiment. Groups of 48 fish were prepared in a 48-well plate according to Joo et al. ^86^ (Fig. 1a) and subjected to different sleep disruption protocols while their locomotor activity was monitored for 24 hours. Visible light of around 2500 lux was switched on during the night ZT14-ZT0 (11 PM - 9 AM) according to different sleep protocols (Fig. S1a). 1) Light: The light was on constantly during the night. 2) Light pulses: The light was switched off for 15 min and then switched on for 45 min. This was repeated 10 times throughout the night. 3) 6h light: The light was switched off for 4 h and then switched on for the remaining 6 h (ZT18-ZT0) of the night. For the control group, the light was switched off during the night. All groups were subjected to light from ZT0-ZT14 (9 AM-11PM) on.

To test sleep rebound, sleep disruption at night was followed by a light period of 3 h between ZT0-ZT3 (9 AM - 12 PM) and a subsequent dark period (Fig. 1f-h, S1e-h).

Custom-written programs allowed subsequent analysis of the fish’s locomotion as described in Joo et al. (2021) ^86^. Activity during the recording period was determined by subtracting subsequent frames from each other and using the number of pixels that changed intensity as a readout. Sleep was defined as an absence of movement for at least one minute.

Due to space constraints in the 48-well plate and to exclude effects of social isolation ^87^, groups of 20 fish were sleep disrupted in a bigger round dish (⌀ 10 cm) when behavioral experiments followed the disturbance. If not stated otherwise, in visual-based decision-making experiments, fish of the AB strain were sleep disrupted using continuous illumination at night (Fig. 2). In olfactory-based decision-making experiments (Fig. 4), fish of the AB strain were sleep disrupted with light pulses.

#### Swim Monitoring

The fish’s swimming was monitored over 24 hours (Fig. 1i-m, Fig. S1 i-l) in the setup described in Bahl and Engert (2020) ^27^. At night, one group of larvae was sleep disrupted using light from below while the other group was monitored over an undisturbed night. Single larvae were placed in 16 petri dishes, 12 cm in diameter (10 mm in height, black rim, transparent bottom) filled with 100 mL filtered fish facility water. Infrared (IR) light from the bottom allowed tracking of the fish in a region of interest with a camera (Grasshopper3-NIR, FLIR Systems) equipped with a zoom lens (Zoom 7000, 18–108 mm, Navitar) and a long-pass filter (R72, Hoya). Python software developed in the Engert laboratory ^27^ allowed the determination of the fish’s position in real-time. Bright LED floodlights (approximately 2500 lux) were mounted below the dishes for sleep disruption, each light illuminating 4 dishes. From ZT7-ZT14 (4 PM – 11 PM) the floodlights were switched on for all fish. For the control group, the lights were switched off during the night ZT14-ZT0(11 PM – 9 AM) while for the sleep-disrupted groups, the lights were switched on during the night either constantly or with light pulses (21 minutes of darkness, 64 minutes of light; repeated 7 times). At ZT0 the light was switched back on for all groups until ZT7 (4 PM). Water in the dishes was refilled at ZT0 (9 AM) due to evaporation over the long time period. During the whole experiment the fish were tracked, and their position and swim bouts were determined.

In Figure 2 l-n, fish underwent sleep disruption for one night as described above, before they were tested in the visual-based decision-making task. Single fish experienced either total darkness for one night, continuous light or a one hour light pulse from ZT15-ZT16 (12 AM to 1AM).

### Vision-based decision-making task (optomotor response)

Decision-making ability was tested using the optomotor response (OMR). Experiments were performed in the setup described in Bahl and Engert (2020) ^27^. In short, larvae were placed in a petri dish while a dot motion kinematogram was projected onto the bottom of the dish. The dots were moving with either 0, 25, 50, or 100 % coherence in one direction perpendicular to the fish in a closed loop. The remaining dots were moving randomly. The fish’s position, and orientation were tracked in real time with a custom software ^27^. Subsequently, different behavioral parameters were analyzed using a custom-written Python script.

Performance was determined as the probability of making a turn in direction with the movement of the visual stimulus.

#### Sleep disruption

Larvae were raised under the regular 14 h light/10 h dark sleep cycle and fed daily at ZT0. On two consecutive days between ZT0 and ZT4 (9 AM - 1 PM), the optomotor response was tested in the free-swimming setup described above with moving dots of 0, 25, 50, and 100 % coherence (Fig. S2m-p), once before and once after sleep disruption. Right before the experiments on both days, fish were placed under a bright LED lamp (1000 lux) to prevent sleep-disrupted animals from catching up on sleep during the day. Each fish underwent 30 trials which combined to a 60-minute experimental period.

#### Melatonin treatment

Larvae were raised under the regular 14 h light/10 h dark sleep cycle. After an initial test in the decision-making task (visual-based decision-making task, optomotor response) in fish water, all larvae were tested in a melatonin solution of varying concentrations (Fig. 3a-e, Fig. S4). Melatonin stock solutions of 0.1 mM or 0.01 mM (for a final concentration of 1 nM) were prepared by adding melatonin (Selleckchem, Catalog No. S1204, 10mM) to DMSO (Electron Microscopy Sciences 13390). To each dish containing 50 ml of fish water, we added one of the following: 50 μl of the 0.1 mM melatonin stock solution (final melatonin concentration of 100 nM; final DMSO concentration of 0.1%), 5 μl of the 0.1 mM melatonin stock (final melatonin concentration of 10 nM; final DMSO concentration of 0.01%), or 5 μl of the 0.01 mM melatonin stock (final melatonin concentration of 1 nM; final DMSO concentration of 0.01%). To exclude effects of DMSO, one initial experiment was done with sibling controls, either treated with 100 nM melatonin or 0.1% DMSO (Fig. S4a) after the first round of OMR testing in fish water. Behavioral testing concentrations were selected based on Ghosh and Rihel (2020) ^39^. Fish were removed from the dishes while melatonin was added and afterwards placed back into their respective petri dishes. After a 20-min incubation, the fish were presented with the same decision-making task. The trial number of the four coherent stimuli amounted to 30 trials per round. Each round lasted 60 min.

In the recovery experiment, the melatonin solution was removed and replaced by fresh water from the fish facility (Fig. S4b). After a recovery period of 20 min a third decision-making task was started. In this experiment, the animals underwent 20 trials of the 4 different stimuli to keep the total trial number for all fish at 60 trials and to prevent reduced bouting due to fatigue.

#### Cortisol treatment

Larvae of the AB strain were raised under the regular 14h light/10 h dark sleep cycle. A hydrocortisone stock solution of 1 mg/ml was prepared by dissolving hydrocortisone (Sigma Aldrich) in fish water. After the initial decision-making task in fish water, larvae were removed from the dish, and 25 μl of the hydrocortisone stock solution (Sigma Aldrich) were added to the dish containing 50 ml fish water for a final concentration of 5 μg/ml cortisol. Larvae were placed back into their respective petri dishes. After a 20-minute incubation, the fish were presented with the same decision-making task (Fig. 5f-i). The trial number of the four coherent stimuli amounted to 30 trials per round. Each round lasted 60 min.

#### High bouter - Low bouter

Controls that underwent the OMR paradigm in previous experiments were analyzed for their bout frequency (bouts/min). Fish with the lowest 25% of bout frequency were used to form a new group of low-bouters and fish with the highest 25% of bout frequency formed the high-bouters. The OMR performance of these two groups was then analyzed separately.

### Olfaction-based decision-making task (Avoidance Assay)

Olfactory avoidance behavior was tracked using the setup described in Herrera et al. (2021) ^34^ (Fig. 4a). In short, a chemical gradient was created by adding chemical scents of cadaverine (Sigma Aldrich), putrescine dihydrochloride (Sigma Aldrich), or sodium chloride (NaCl, VWR) into a 3% low melting point (LMP) agarose (AquaPor LM GTAC) solution before solidification and placing agarose gels at either end of a shallow four laned dish. One gel consisted of pure agarose while the other contained the chemical sent. Cadaverine concentrations of 0 mM, 1 mM, 2.5 mM, and 5 mM were tested while a putrescine concentration of 10 mM was tested (Fig. 4). A salt concentration of 100 mM was tested (Fig. 4). The lanes were filled with filtered fish water, allowing for the diffusion of the chemical in the water to form a scent gradient. In the case of cortisol treatment described below, the lanes were filled with cortisol solution of the respective concentration.

Immediately, after water was added, four larvae were placed into the middle of each lane to avoid biasing their side preference. An IR camera positioned above the dish was tracking fish movement and location. An LED light positioned at the side of the setup illuminated the lanes for the fish and prevented daytime sleep. Using a LabVIEW program created by Hererra et al. ^34^, fish were tracked over a 30-min period. The location (left or right) of the chemical-containing agar was randomized and noted in the experiment file for later analysis.

Preference index (PI), bout rate, bout probability and turning angles were analyzed using custom-written code. The script calculated the PI based on the position of the fish along the axis perpendicular to the agar pads. The PI is here defined as the difference between the time fish spend in the half of the dish near the chemically treated agarose and the time spent on the plain agarose side divided by the total time.

PI = (time near scent- time far from scent) / (total time)

Every fish’s average bout rate difference was calculated by subtracting the bout rate per minute within 2 cm of the plain agarose from the bout rate within 2 cm of the odor source.

Bout rate difference = Average bout rate near scent (bouts/min) - Average bout rate far from scent (bouts/min)

The bout count probability was determined as the proportion of bouts that were found within a bin. Each bin made up 1 mm of the lanes in which the odor concentration gradient was formed, effectively describing a relative local odor concentration.

Heading angles were calculated as described in Herrera et al. (2021) ^34^. In brief, high variances in heading angles were identified from segmented bouts. The bouts were then separated into by whether the fish moves towards the scented agarose pad or away from it (at least 0.2 mm). Fish that were not tracked or did not move for more than 70% of the time were removed from the analysis. Bouts that occurred within 50 ms of another bout were removed from the analysis.

#### In sleep-disrupted fish

To test sleep-disrupted fish in the olfactory-based decision-making task, sibling larvae were randomly separated into control and sleep-disrupted groups, consisting of approximately 20 fish. Overnight, they were subjected to either control darkness or a light pulse protocol (Sleep Monitoring). Due to limitations of the setup, for the avoidance behavior using putrescine, some fish were subjected to 10 light pulses consisting of 15 minutes darkness and 45 minutes light, and others were subjected to 7 light pulses consisting of 21 minutes of darkness and 64 minutes of light. At ZT0 (9 AM), fish were collected and fed paramecia. They were then left under an LED light for 30 min before beginning the experiments. A group of four fish were tested at a time, alternating control and sleep-disrupted groups. Olfactory avoidance was tested with different cadaverine concentrations as described above and with putrescine and salt (Fig. 4, S6).

#### In cortisol-treated fish

Larvae were randomly separated into control and experimental groups consisting of approximately 20 fish and kept in darkness overnight. At ZT0, fish were fed in the morning and placed under a bright LED lamp for 30 min. Subsequently, fish were tested in the olfactory-based decision-making task alternating between control and experimental. Experimental fish were placed in either a 5 ng/ml, 100 ng/ml, or 5 μg/ml hydrocortisone solution 15 min prior to experimentation and were tested in the same concentration (Fig. 5, S7). Control fish were tested in fish water.

#### In starved fish

To simulate the increased metabolic demand larvae underwent a partial or a complete starvation protocol (Fig. S7). For partial starvation, the starvation group was not fed on the day prior to the behavioral experiment, while the control group was fed at ZT0 (9 AM). On the day of the experiment, both groups were fed with paramecia at ZT0 (9 AM) and placed under a bright LED light for 30 min before being subjected to the olfactory-based decision-making task. For complete starvation, both control and starved groups were fed at ZT0 (9 AM) and ZT9 (6 PM) with plenty of paramecia on the day prior to the experiment. Only the control group was fed in the morning of the experiment (ZT0), while the starvation group was moved to a fresh dish free of paramecia. Both groups were placed under a bright LED light for 3 h after feeding and subsequently performed the olfactory-based decision-making task.

### Cortisol luminescence assay

Cortisol concentration was quantified using a luminescence assay as described in Rennekamp et al. (2016) ^88^. Cortisol was measured in sleep-disrupted fish, larvae treated with cortisol, and fish stressed with an electric shock, all with respective controls (Fig. 5a, Fig. S7f). Sibling fish of the AB strain were raised under a 14h light/10 h dark cycle in a petri dish (15 cm in diameter) in filtered fish facility water at 28°C. Samples for cortisol quantification were collected following one of three protocols: 1) Larvae were randomly separated into groups of 30 and sleep disrupted with continuous light or light pulses as described above. The control group was kept in darkness during the night. Fish were removed from the setup promptly at ZT0 (9 AM) on the following day and fed while being kept under a bright LED lamp. After resting for 30 min, the dishes were placed on ice. Half the water from each dish was removed and replaced with ice water. This was repeated two more times until the fish ceased swimming. Fish were transferred to prechilled 1.5 mL tubes, spun down for 5 s, and placed back on ice. The water was removed from the tubes and the fish were frozen in the tubes using a dry ice ethanol bath. Samples were stored at −20°C until further preparation. 2) At ZT0, all fish were fed and after 30 min a group of 30 fish was placed one by one in a square tank and shocked once with 5V while the control group was just placed in the tank and removed again after 20 s. After 15 min, both tanks were filled with ice water to render the fish immobile and the larvae were frozen as described above. 3) At ZT0, siblings were fed and after 30 min, placed in dishes containing one of the following hydrocortisone (Sigma Aldrich) concentrations: 5 ng/ml, 100 ng/ml, 5 μg/ml, or 10 μg/ml. The control group was placed in filtered fish water. After 30 min, all dishes were filled with ice water and fish were frozen at −20°C. After all samples were collected, cortisol was extracted following the protocol by Rennekamp et al. (2016) ^88^ using the Cortisol Saliva Luminescence Immunoassay Kit (IBL International). Luminescence was read out using the Biotek Neo2 Luminescence Reader in Harvard’s Bauer Core facility. The internal cortisol concentrations were determined using a standard curve and interpolation with GraphPad Prism. To normalize the data, the mean control values were subtracted from the respective treatments.

### per3:luciferase assay

The per3:Luc circadian rhythm bioluminescence assay (Fig. 2f - k) was performed as described in Suppermpool et al. (2023)^19^. per3:Luc larvae (*Tg(per3:luc)* in which the promoter of *period 3* gene drives expression of luciferase ^89^ were entrained on a 14h light/10 h dark cycle until 5 dpf and then placed in groups of 50 in ⌀10 cm petri dishes in water containing 0.5 mM beetle luciferin (Promega) in 2500 Lux illumination. From ZT14 (11 PM) on 5 dpf these larvae experienced one of three different lighting paradigms: the control group continued on the 14h light/10 h dark cycle; the constant light sleep disruption group were exposed to constant 2500 lux light throughout the night; the ZT15-16 1h Light Pulse group were exposed to 1h at 0 lux, followed by 1h at 2500 Lux (ZT15-16, 12 AM - 1 AM), and then 0 lux for a further 8 hours. All three groups were then exposed to approximately 1 h of 2500 lux illumination from control ZT0 (09 AM) on 6 dpf, before larvae were transferred individually into a white-walled 96-well plate (Greiner Bio-One) and sealed with oxygen-permeable plate-seal (Applied Biosystems). These plates were maintained at 28 ° C in constant darkness and bioluminescence photon counts were sampled every 20 min for >130 h using a TopCount NXT scintillation counter (Packard) to measure the expression of luciferase under the control of the per3 promoter. Circadian period and phase were analyzed using BioDare2 (biodare2.ed.ac.uk; Zieliński et al. 2022 ^90^) and is BioDare2 experiment number 29533.

### Statistical analysis

Statistical analysis was performed in Python and MATLAB using parametric statistical models. Specifically, for between-subject experiments (Sleep Disruption), we used two-way mixed ANOVAs followed by Tukey’s HSD post hoc tests and two-sided independent t-tests. For within-subject experiments (Treatments) we used two-way repeated measures ANOVAs followed by Tukey’s HSD post hoc tests and two-sided t-tests for paired samples. Wilcoxon rank sum tests were used for unequal subject sizes. Parametric distributions can be assumed due to the large sample sizes in our experiments. P-values are reported in the figure legends. Spearman’s rank correlation coefficient was calculated using the Python package scipy.stats.

### Computational Modeling

Algorithmic modeling using the leaky integrator model in freely swimming larvae was performed as described previously in (Harpaz et al 2021) ^33^. In brief, we implemented an integrator variable (x) that accumulates Gaussian noise () and coherence-dependent motion drift (c). The integrator is leaky, such that it slowly decayed to zero with a time constant () without an external stimulus:

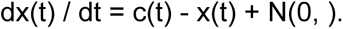

N(= 0,) is a random Gaussian variable with a mean of = 0 and a standard deviation. We simulated this stochastic differential equation using a forward Euler approach with dt = 0.01 s. At each time step, we compared the value of the variable x with the threshold T. When -T < x < T, we considered the variable to be in the forward swimming regime. When x >= T or x <= -T, the model fish swam to the right or left, respectively. Swimming was initiated stochastically, with two different probabilities (pbelow and pabove), depending on whether x was below or above the threshold, respectively. Furthermore, right and left turns were drawn from Gaussian distributions with N( = 22, = 25) and, respectively. Forward swims were drawn from N(= 0, = 5). These distribution choices approximated the heading angle change distributions that we observed in our experiments (Fig. 2c).

Following our recent work (Harpaz et al. 2021) ^33^, we chose the following baseline parameters: =0.8, = 10, T = 1, pbelow= 0.008, pabove=0.04. These parameters produced swim rates and performances that capture the measured behavior well. To test the impact of specific parameters, we systematically adjusted T,, and pbelow, while keeping the other parameters at their respective baseline values. T was varied between [1, 1.2, 1.4, 1.5], between [0.8, 1.12, 1.44, 1.6], and p_below and p_above between [0.040, 0.024, 0.008, 0.0000]. At each swim, the position was updated with 0.5 cm in the direction of the swim. The input evidence x was directly corresponding to the dot-motion coherence open-loop simulations, whereas for the closed-loop it was computed as the coherence times the sine of the relative angle between the motion and the simulated fish. Model data was then saved in the form of swim events, using a similar data format as in our experiments, allowing us to use the same statistical framework for analyzing events and plotting the data.

## Data Availability

The data collected in this study will be made available through Harvard University’s data repository system.

## Code Availability

The software used to analyze the data generated in this study was developed in Python (v3.10), MATLAB (R2022b), and Jupyterlab ^91^. The analysis code will be made available on GitHub.

## Acknowledgments

Special thanks to Jessica Miller and Karen Hurley for their guidance with fish care. We extend our thanks to Roy Harpaz for his valuable discussions and insights into statistical testing. Appreciation is also extended to Constance Richter and Terzah Hill for their assistance with cortisol measurements. Lastly, we express our thanks to all members of the Engert lab for their support and guidance throughout the project.

A.B. and M.C. received funding through the Emmy Noether Program (BA 5923/1-1), the Excellence Strategy (EXC 2117-422037984), and the Zukunftskolleg Konstanz.

## Author contributions

H.Z. designed the experiments after initial discussions with J.R., D.G.L., and W.J.. P.P., N.O., K.K., W.J. and D.G.L. performed experiments. P.P., N.O., K.K., K.J.H., D.G.L. and H.Z. analyzed the data. A.B. and M.C. conducted computational modeling. P.P. and H.Z. wrote the paper with revisions from K.J.H., W.J., A.B., F.E., D.G.L. and J.R. F.E. and H.Z. supervised the project.

## REFERENCES

1. Goel, N., Rao, H., Durmer, J. S. & Dinges, D. F. Neurocognitive Consequences of Sleep Deprivation. Semin. Neurol. 29, 320 (2009).

2. Harrison, Y. & Horne, J. A. One Night of Sleep Loss Impairs Innovative Thinking and Flexible Decision Making. Organ. Behav. Hum. Decis. Process. 78, 128–145 (1999).

3. Killgore, W. D. S., Balkin, T. J. & Wesensten, N. J. Impaired Decision Making Following 49 h of Sleep Deprivation. 7–13 (2006).

4. Rodgers, D., et al. Sleep Deprivation: Effects on Work Capacity, Self-Paced Walking, Contractile Properties and Perceived Exertion. Sleep vol. 18 30–38 (1995).

5. Salfi, F. et al. Effects of Total and Partial Sleep Deprivation on Reflection Impulsivity and Risk-Taking in Deliberative Decision-Making. Nat. Sci. Sleep 2020, 12–309 (2020).

6. Whitney, P., Hinson, J. M., Jackson, M. L. & Van Dongen, H. P. A. Feedback Blunting: Total Sleep Deprivation Impairs Decision Making that Requires Updating Based on Feedback. Sleep 38, 745–754 (2015).

7. Williamson, A. M., Feyer, A.-M., Wales, S. & M Williamson, A. A. Moderate sleep deprivation produces impairments in cognitive and motor performance equivalent to legally prescribed levels of alcohol intoxication. Occup Env. Med 57, 649–655 (2000).

8. Prober, D. A., Rihel, J., Onah, A. A., Sung, R. J. & Schier, A. F. Hypocretin/Orexin Overexpression Induces An Insomnia-Like Phenotype in Zebrafish. J. Neurosci. 26, 13400–13410 (2006).

9. Zhdanova, I. V., Wang, S. Y., Leclair, O. U. & Danilova, N. P. Melatonin promotes sleep-like state in zebrafish. Brain Res. 903, 263–268 (2001).

10. Gandhi, A. V., Mosser, E. A., Oikonomou, G. & Prober, D. A. Melatonin Is Required for the Circadian Regulation of Sleep. Neuron 85, 1193–1199 (2015).

11. Reichert, S., Pavón Arocas, O. & Rihel, J. The Neuropeptide Galanin Is Required for Homeostatic Rebound Sleep following Increased Neuronal Activity. Neuron 104, 370–384.e5 (2019).

12. Oikonomou, G. et al. The Serotonergic Raphe Promote Sleep in Zebrafish and Mice. Neuron 103, 686–701.e8 (2019).

13. Mommsen, T. P., Vijayan, M. M. & Moon, T. W. Cortisol in teleosts: Dynamics, mechanisms of action, and metabolic regulation. Rev. Fish Biol. Fish. 9, 211–268 (1999).

14. Aluru, N. & Vijayan, M. M. Stress transcriptomics in fish: A role for genomic cortisol signaling. Gen. Comp. Endocrinol. 164, 142–150 (2009).

15. Pennisi, E. The simplest of slumbers. Science 374, 526–529 (2021).

16. Zimmerman, J. E., Naidoo, N., Raizen, D. M. & Pack, A. I. Conservation of sleep: insights from non-mammalian model systems. Trends Neurosci. 31, 371–376 (2008).

17. Franken, P. & Dijk, D.-J. Sleep and circadian rhythmicity as entangled processes serving homeostasis. Nat. Rev. Neurosci. 25, 43–59 (2024).

18. Elbaz, I., Yelin-Bekerman, L., Nicenboim, J., Vatine, G. & Appelbaum, L. Genetic Ablation of Hypocretin Neurons Alters Behavioral State Transitions in Zebrafish. J. Neurosci. 32, 12961–12972 (2012).

19. Suppermpool, A., Lyons, D. G., Broom, E. & Rihel, J. Sleep pressure modulates single-neuron synapse dynamics in zebrafish. 2023.08.30.555615 Preprint at 10.1101/2023.08.30.555615 (2023).

20. Leung, L. C. et al. Neural signatures of sleep in zebrafish. Nature 571, 198–204 (2019).

21. Aho, V. et al. Homeostatic response to sleep/rest deprivation by constant water flow in larval zebrafish in both dark and light conditions Sleep homeostasis in zebrafish. J Sleep Res 26, 394–400 (2017).

22. Yokogawa, T. et al. Characterization of Sleep in Zebrafish and Insomnia in Hypocretin Receptor Mutants. PLOS Biol. 5, e277 (2007).

23. Silva, R. F. O., Pinho, B. R., Santos, M. M. & Oliveira, J. M. A. Disruptions of circadian rhythms, sleep, and stress responses in zebrafish: New infrared-based activity monitoring assays for toxicity assessment. Chemosphere 305, 135449 (2022).

24. Zada, D. et al. Parp1 promotes sleep, which enhances DNA repair in neurons. Mol. Cell 81, 4979–4993.e7 (2021).

25. Oikonomou, G. & Prober, D. A. Attacking sleep from a new angle: contributions from zebrafish. Curr. Opin. Neurobiol. 44, 80–88 (2017).

26. Rihel, J., Prober, D. A. & Schier, A. F. *Monitoring Sleep and Arousal in Zebrafish*. Methods in Cell Biology vol. 100 (Academic Press, 2010).

27. Bahl, A. & Engert, F. Neural circuits for evidence accumulation and decision making in larval zebrafish. Nat. Neurosci. 23, 94–102 (2020).

28. Cherng, B.-W., Islam, T., Torigoe, M., Tsuboi, T. & Okamoto, H. The Dorsal Lateral Habenula-Interpeduncular Nucleus Pathway Is Essential for Left-Right-Dependent Decision Making in Zebrafish. Cell Rep. 32, 108143 (2020).

29. Barker, A. J. & Baier, H. Sensorimotor Decision Making in the Zebrafish Tectum. Curr. Biol. 25, 2804–2814 (2015).

30. Neuhauss, S. C. F. et al. Genetic Disorders of Vision Revealed by a Behavioral Screen of 400 Essential Loci in Zebrafish. J. Neurosci. 19, 8603–8615 (1999).

31. Kist, A. M. & Portugues, R. Optomotor Swimming in Larval Zebrafish Is Driven by Global Whole-Field Visual Motion and Local Light-Dark Transitions. Cell Rep. 29, 659–670.e3 (2019).

32. Götz, K. G., Hengstenberg, B. & Biesinger, R. Optomotor control of wing beat and body posture in drosophila. Biol. Cybern. 35, 101–112 (1979).

33. Harpaz, R. et al. Collective behavior emerges from genetically controlled simple behavioral motifs in zebrafish. Sci. Adv. 7, (2021).

34. Herrera, K. J., Panier, T., Guggiana-Nilo, D. & Engert, F. Larval Zebrafish Use Olfactory Detection of Sodium and Chloride to Avoid Salt Water. Curr. Biol. 31, 782–793.e3 (2021).

35. Koide, T., Yabuki, Y. & Yoshihara, Y. Terminal Nerve GnRH3 Neurons Mediate Slow Avoidance of Carbon Dioxide in Larval Zebrafish. Cell Rep. 22, 1115–1123 (2018).

36. Sy, S. K. H. et al. An optofluidic platform for interrogating chemosensory behavior and brainwide neural representation in larval zebrafish. Nat. Commun. 14, 227 (2023).

37. Hussain, A. et al. High-affinity olfactory receptor for the death-associated odor cadaverine. Proc. Natl. Acad. Sci. 110, 19579–19584 (2013).

38. Krishnan, K., et al. Attentional Switching in Larval Zebrafish: The Attentive Leaky Integrator. (2023). doi:10.21203/rs.3.rs-3486824/v1.

39. Ghosh, M. & Rihel, J. Hierarchical Compression Reveals Sub-Second to Day-Long Structure in Larval Zebrafish Behavior. eNeuro 7, 1–21 (2020).

40. Zhdanova, I. V. Sleep and its regulation in zebrafish. Rev. Neurosci. 22, 27–36 (2011).

41. Everson, C. A. & Wehr, T. A. Nutritional and metabolic adaptations to prolonged sleep deprivation in the rat. Am. J. Physiol.-Regul. Integr. Comp. Physiol. 264, R376–R387 (1993).

42. Greer, S. M., Goldstein, A. N. & Walker, M. P. The impact of sleep deprivation on food desire in the human brain. Nat. Commun. 4, 2259 (2013).

43. Calvin, A. D. et al. Effects of Experimental Sleep Restriction on Caloric Intake and Activity Energy Expenditure. Chest 144, 79–86 (2013).

44. Jung, C. M. et al. Energy expenditure during sleep, sleep deprivation and sleep following sleep deprivation in adult humans. J. Physiol. 589, 235–244 (2011).

45. Markwald, R. R. et al. Impact of insufficient sleep on total daily energy expenditure, food intake, and weight gain. Proc. Natl. Acad. Sci. 110, 5695–5700 (2013).

46. Skajaa, K., Fernö, A. & Folkvord, A. Swimming, feeding and predator avoidance in cod larvae (Gadus morhua L.): trade-offs between hunger and predation risk.

47. Scharf, I. The multifaceted effects of starvation on arthropod behaviour. Anim. Behav. 119, 37–48 (2016).

48. Hoenen, M., Wolf, O. T. & Pause, B. M. The Impact of Stress on Odor Perception. Perception 46, 366–376 (2017).

49. Pause, B. M., Sojka, B., Krauel, K., Fehm-Wolfsdorf, G. & Ferstl, R. Olfactory information processing during the course of the menstrual cycle. Biol. Psychol. 44, 31–54 (1996).

50. Best, C. & Vijayan, M. M. Cortisol elevation post-hatch affects behavioural performance in zebrafish larvae. Gen. Comp. Endocrinol. 257, 220–226 (2018).

51. Zhdanova, I. V. Sleep in Zebrafish. Zebrafish 3, 215–226 (2006).

52. Inostroza, M., Binder, S. & Born, J. Sleep-dependency of episodic-like memory consolidation in rats. Behav. Brain Res. 237, 15–22 (2013).

53. Yang, R.-H. et al. Paradoxical sleep deprivation impairs spatial learning and affects membrane excitability and mitochondrial protein in the hippocampus. Brain Res. 1230, 224–232 (2008).

54. Colavito, V. et al. Experimental sleep deprivation as a tool to test memory deficits in rodents. Front. Syst. Neurosci. 7, 106 (2013).

55. Weger, B. D. et al. The Light Responsive Transcriptome of the Zebrafish: Function and Regulation. PLoS ONE 6, e17080 (2011).

56. Pittaras, E. et al. Mouse Gambling Task reveals differential effects of acute sleep debt on decision-making and associated neurochemical changes. Sleep 41, zsy168 (2018).

57. Seugnet, L., Suzuki, Y., Donlea, J. M., Gottschalk, L. & Shaw, P. J. Sleep deprivation during early-adult development results in long-lasting learning deficits in adult Drosophila. Sleep 34, 137–146 (2011).

58. Seugnet, L., Galvin, J. E., Suzuki, Y., Gottschalk, L. & Shaw, P. J. Persistent Short-Term Memory Defects Following Sleep Deprivation in a Drosophila Model of Parkinson Disease. Sleep 32, 984 (2009).

59. Pinheiro-da-Silva, J., Silva, P. F., Nogueira, M. B. & Luchiari, A. C. Sleep deprivation effects on object discrimination task in zebrafish (Danio rerio). Anim. Cogn. 20, 159–169 (2017).

60. Aguiar, S. A. & Barela, J. A. Sleep deprivation affects sensorimotor coupling in postural control of young adults. Neurosci. Lett. 574, 47–52 (2014).

61. Wang, M. Y., Brennan, C. H., Lachlan, R. F. & Chittka, L. Speed-accuracy trade-offs and individually consistent decision making by individuals and dyads of zebrafish in a colour discrimination task. Anim. Behav. 103, 277–283 (2015).

62. Burns, J. G. Impulsive bees forage better: the advantage of quick, sometimes inaccurate foraging decisions. Anim. Behav. 70, e1–e5 (2005).

63. Chittka, L., Dyer, A. G., Bock, F. & Dornhaus, A. Bees trade off foraging speed for accuracy. Nature 424, 388–388 (2003).

64. Burns, J. G. & Dyer, A. G. Diversity of speed-accuracy strategies benefits social insects. Curr. Biol. CB 18, R953–954 (2008).

65. Franks, N. R., Dornhaus, A., Fitzsimmons, J. P. & Stevens, M. Speed versus accuracy in collective decision making. Proc. R. Soc. Lond. B Biol. Sci. 270, 2457–2463 (2003).

66. Kulahci, I. G., Dornhaus, A. & Papaj, D. R. Multimodal signals enhance decision making in foraging bumble-bees. Proc. R. Soc. B Biol. Sci. 275, 797 (2008).

67. Ducatez, S., Audet, J. N. & Lefebvre, L. Problem-solving and learning in Carib grackles: individuals show a consistent speed–accuracy trade-off. Anim. Cogn. 18, 485–496 (2015).

68. Rinberg, D., Koulakov, A. & Gelperin, A. Speed-Accuracy Tradeoff in Olfaction. Neuron 51, 351–358 (2006).

69. Lim, J. & Dinges, D. F. Sleep deprivation and vigilant attention. Ann. N. Y. Acad. Sci. 1129, 305–322 (2008).

70. Uchida, N. & Mainen, Z. F. Speed and accuracy of olfactory discrimination in the rat. Nat. Neurosci. 6, 1224–1229 (2003).

71. Killgore, W. D. S. & McBride, S. A. Odor identification accuracy declines following 24 h of sleep deprivation. J. Sleep Res. 15, 111–116 (2006).

72. Mcbride, S. A., Balkin, T. J., Kamimori, G. H. & Killgore, W. D. s. Olfactory Decrements as a Function of Two Nights of Sleep Deprivation. J. Sens. Stud. 21, 456–463 (2006).

73. Frontiers | On the state-dependent nature of odor perception. https://www.frontiersin.org/journals/neuroscience/articles/10.3389/fnins.2022.964742/full.

74. Goldstein, A. N. et al. Tired and apprehensive: anxiety amplifies the impact of sleep loss on aversive brain anticipation. J. Neurosci. Off. J. Soc. Neurosci. 33, 10607–10615 (2013).

75. Yoo, S.-S., Gujar, N., Hu, P., Jolesz, F. A. & Walker, M. P. The human emotional brain without sleep — a prefrontal amygdala disconnect. Curr. Biol. 17, R877–R878 (2007).

76. Goldstein-Piekarski, A. N., Greer, S. M., Saletin, J. M. & Walker, M. P. Sleep Deprivation Impairs the Human Central and Peripheral Nervous System Discrimination of Social Threat. J. Neurosci. Off. J. Soc. Neurosci. 35, 10135–10145 (2015).

77. Uchida, N., Kepecs, A. & Mainen, Z. F. Seeing at a glance, smelling in a whiff: rapid forms of perceptual decision making. Nat. Rev. Neurosci. 7, 485–491 (2006).

78. Uchida, N. & Mainen, Z. F. Speed and accuracy of olfactory discrimination in the rat. Nat. Neurosci. 6, 1224–1229 (2003).

79. Hoenen, M., Wolf, O. T. & Pause, B. M. The Impact of Stress on Odor Perception. Perception 46, 366–376 (2017).

80. Sigurgeirsson, B. et al. Sleep–wake dynamics under extended light and extended dark conditions in adult zebrafish. Behav. Brain Res. 256, 377–390 (2013).

81. Leproult, R., Copinschi, G., Buxton, O. & Van Cauter, E. Sleep Loss Results in an Elevation of Cortisol Levels the Next Evening. Sleep 20, 865–870 (1997).

82. Redwine, L., Hauger, R. L., Gillin, J. C. & Irwin, M. Effects of Sleep and Sleep Deprivation on Interleukin-6, Growth Hormone, Cortisol, and Melatonin Levels in Humans. J. Clin. Endocrinol. Metab. 85, 3597–3603 (2000).

83. Wright, K. P. et al. Influence of sleep deprivation and circadian misalignment on cortisol, inflammatory markers, and cytokine balance. Brain. Behav. Immun. 47, 24–34 (2015).

84. Salín-Pascual, R. J. et al. The effect of total sleep deprivation on plasma melatonin and cortisol in healthy human volunteers. Sleep 11, 362–369 (1988).

85. Pause, B. M., Sojka, B., Krauel, K., Fehm-Wolfsdorf, G. & Ferstl, R. Olfactory information processing during the course of the menstrual cycle. Biol. Psychol. 44, 31–54 (1996).

86. Joo, W., Vivian, M. D., Graham, B. J., Soucy, E. R. & Thyme, S. B. A Customizable Low-Cost System for Massively Parallel Zebrafish Behavioral Phenotyping. Front. Behav. Neurosci. 14, (2021).

87. Wee, C. L. et al. Social isolation modulates appetite and defensive behavior via a common oxytocinergic circuit in larval zebrafish 2 3. doi:10.1101/2020.02.19.956854.

88. Rennekamp, A. J. et al. σ1 receptor ligands control a switch between passive and active threat responses. Nat. Chem. Biol. 12, 552–558 (2016).

89. Kaneko, M. & Cahill, G. M. Light-Dependent Development of Circadian Gene Expression in Transgenic Zebrafish. PLOS Biol. 3, e34 (2005).

90. Zieliński, T., Hay, J. & Millar, A. J. Correction to: Period Estimation and Rhythm Detection in Timeseries Data Using BioDare2, the Free, Online, Community Resource. in Plant Circadian Networks: Methods and Protocols (eds. Staiger, D., Davis, S. & Davis, A. M.) C1–C1 (Springer US, New York, NY, 2022). doi:10.1007/978-1-0716-1912-4_19.

91. Kluyver, T. et al. Jupyter Notebooks—a publishing format for reproducible computational workflows. Position. Power Acad. Publ. Play. Agents Agendas - Proc. 20th Int. Conf. Electron. Publ. ELPUB 2016 87–90 (2016) doi:10.3233/978-1-61499-649-1-87.

