## Supplementary Figures for "Sleep Disruption Improves Performance in Simple Olfactory and Visual Decision-Making Tasks"

### SUPPLEMENTARY MATERIAL

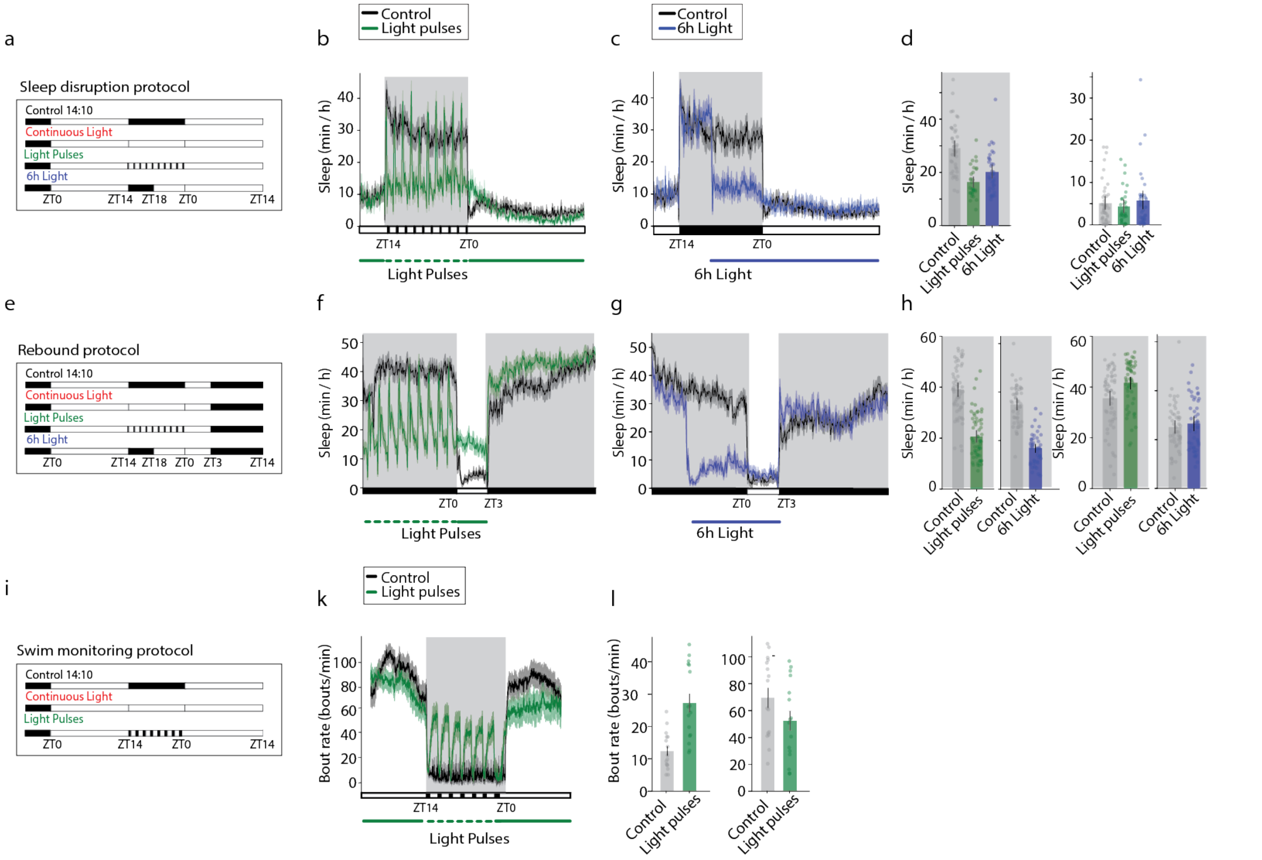


**Figure S1: Light at night prevented sleep and decreased locomotion. a:** Sleep protocols for the recording period. The control group experienced 10 h darkness during the night ZT14-ZT0 (11 PM - 9 AM) while the sibling groups were disrupted with either continuous light, light pulses (15 minutes of darkness, 45 minutes of light; repeated 10 times) or six hours of premature light ZT18-ZT0 (3 AM - 9 AM) during the night. At the start of the following day (ZT0) all lights were turned on. **b:** Light pulses at night disrupted sleep. Control animals that experienced constant darkness slept through the night. $n_{ctrll}$ = 43, $n_{lp}$ = 31. **c:** Premature light of 6h reduced sleep as soon as the light was switched on. $n_{control}$ = 43, $n_{6h}$ = 36. **d:** Mean time spent sleeping during night and day. Data from (b) and (c). Light pulses and 6h of light at night reduced sleep (independent samples t-test: *control vs light pulses*: $p < 0.001$, $t = 7.19$, $df = 70.58$; *control vs 6h light*: $p < 0.001$, $t = 4.61$, $df = 78$; $n_{ctrl}$ = 43, $n_{lp}$ = 31, $n_{6h}$ = 36). **e:** Rebound protocol for the recording period. Sleep disruption as in (a) was followed by 3 hours of light before the light was switched off for all groups ZT3 (12 PM). **f:** Fish disrupted with light pulses had increased sleep in the following daytime darkness, especially in the beginning of the dark period. $n_{ctrl}$=58, $n_{lp}$= 59. **g:** Fish disrupted with 6h of premature light showed marginally increased sleep in the beginning of the dark period. $n_{ctrl}$ = 48, $n_{6h}$ = 59. **h:** Mean time spent sleeping during the night and the following daytime darkness. Data from (f) and (g). Sleep disruption with light pulses and 6h light decreased sleep during disturbance (independent samples t-test: *control vs light pulses*: $p < 0.001$, $t = 10.56$, $df = 113.5$; control vs 6h light: $p < 0.001$, $t = 12.13$, $df = 87.57$) and increased sleep in a following dark period (independent samples t-test: control vs light pulses: $p = 0.002$, $t = -3.16$, $df = 108.2$; control vs 6h light: $p = 0.5$, $t = -0.67$, $df = 93.72$). **i:** Swim monitoring protocol for the recording period. The control group experienced regular darkness for 10 h during the night while the sibling groups were disrupted with either continuous light or light pulses (21 minutes of darkness, 64 min of light; repeated 7 times). **k:** Sleep disruption with light pulses at night increased the bout rate during lights-on periods and reduced the bout rate on the following day. n = 16/group. **l**: Mean bout rate from (k). The bout rate of fish who's sleep was disrupted with light was increased at night (independent samples t-test: p < 0.001, t = -

5.91, df = 18.36, n = 16/group) and decreased on the following day (independent samples t-test: $p = 0.08$, $t = 1.8$, $df = 29.48$). Graphs represent mean ±SEM with the responses of individual fish overlaid in circles.

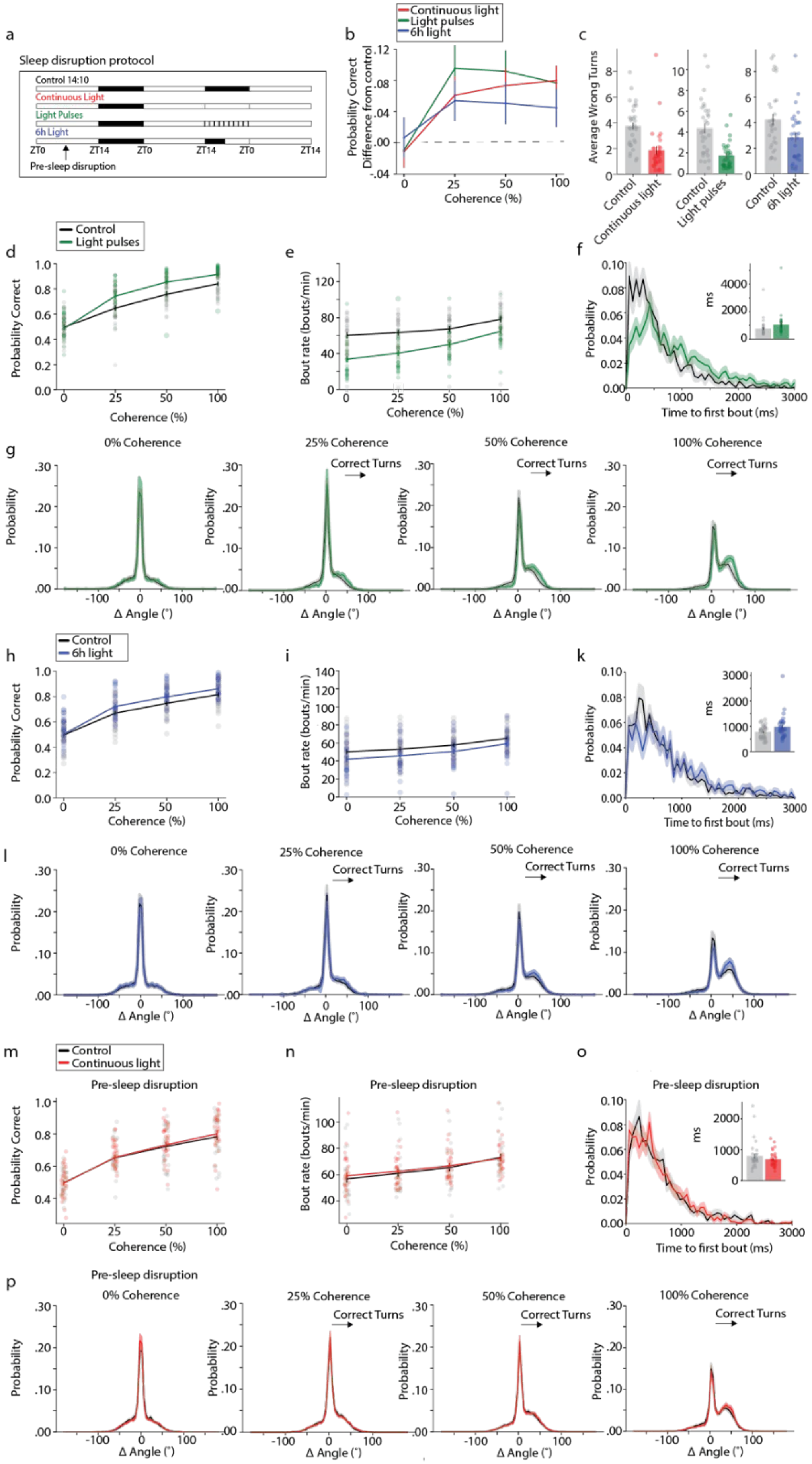
a
Sleep disruption protocol
Control 14:10
Continuous Light
Light Pulses
6h Light
ZT0
ZT14
ZT0
ZT14
ZT0
ZT14
Pre-sleep disruption
b
Continuous light
Light pulses
6h light
Probability Correct Difference from control
Coherence (%)
c
Average Wrong Turns
Control
Continuous light
Control
Light pulses
Control
6h light
d
Control
Light pulses
Probability Correct
Coherence (%)
e
Bout rate (bouts/min)
Coherence (%)
f
Probability
Time to first bout (ms)
ms
g
0% Coherence
25% Coherence
50% Coherence
100% Coherence
Correct Turns
Probability
Δ Angle (°)
h
Control
6h light
Probability Correct
Coherence (%)
i
Bout rate (bouts/min)
Coherence (%)
k
Probability
Time to first bout (ms)
ms
l
0% Coherence
25% Coherence
50% Coherence
100% Coherence
Correct Turns
Probability
Δ Angle (°)
m
Control
Continuous light
Pre-sleep disruption
Probability Correct
Coherence (%)
n
Pre-sleep disruption
Bout rate (bouts/min)
Coherence (%)
o
Pre-sleep disruption
Probability
Time to first bout (ms)
ms
p
Pre-sleep disruption
0% Coherence
25% Coherence
50% Coherence
100% Coherence
Correct Turns
Probability
Δ Angle (°)

**Figure S2: OMR performance increase was only observed after sleep disruption. a:** Sleep disruption protocol. All fish underwent a regular night with 10 h of darkness. On the following day, all animals performed the optomotor response (OMR) task. Subsequently, fish were split into groups. The control group experienced another regular night, while the sleep disruption groups were subjected to either constant light, light pulses or 6 h of light. Then, all animals performed the OMR task again. If not stated otherwise, the following plots depict the data recorded following the disruption. **b:** All sleep disruption protocols caused fish to perform better than the control. n = 32/group. **c:** The total number of wrong turns across all coherences was lower in sleep-disrupted animals than in control groups (independent samples t-test: Light: $p < 0.001$, t = 4.38, df = 61, n = 32/group; Light pulses: $p < 0.001$, t = 5.01, df = 61, n = 32/ group; 6h light: p = 0.017, t = 2.46, df = 61, n = 32/group). **d-g:** Optomotor response of fish sleep-disrupted with light pulses. **d:** Fish disrupted with light pulses showed increased performance in the visual decision-making task (mixed ANOVA: $p_{group} < 0.001$, F(1,30) = 18.31; Tukey's HSD: $p_{0\%} = 0.46$, $p_{25\%} < 0.001$, $p_{50\%} < 0.001$, $p_{100\%} < 0.001$, n= 32/group). **e:** The bout rate was decreased (mixed ANOVA: $p_{grp} < 0.001$, F(1,30) = 15.7; Tukey's HSD: $p_{0\%} < 0.001$, $p_{25\%} < 0.001$, $p_{50\%} < 0.001$, $p_{100\%} = 0.003$, n= 32/group). **f:** Sleep disruption with light pulses elongated the reaction time compared to control fish (independent samples t-test: p = 0.12, t = -1.58, df = 61, n = 32/group). **g:** The turning angles into the correct direction increased with stimulus coherence in sleep-disrupted fish. **h-l:** Optomotor response of fish disrupted with 6 h of premature light. **h:** Sleep disruption with 6 h of premature light increased the larvae's OMR performance for coherent dot motion compared to control fish (mixed ANOVA: $p_{group} = 0.009$, F(1,30) = 7.72; Tukey's HSD: $p_{0\%} = 0.46$, $p_{25\%} = 0.011$, $p_{50\%} = 0.002$, $p_{100\%} = 0.041$, n= 32/group). **i:** Compared to control fish, the larvae's bout rate was slightly decreased after sleep disruption with 6 h of light (mixed ANOVA: $p_{grp} = 0.055$, F(1,30) = 4.0, n= 32/ group). **k:** The time to first bout for a 100% coherent stimulus was increased in sleep-disrupted fish compared to the control group. **l:** For high coherences, sleep-disrupted larvae exhibited increased turning angles in direction of the visual stimulus (independent samples t-test: p = 0.033, t = -2.18, df = 61, n = 32/group). **m - p:** Visual decision-making task tested before sleep disruption. n = 32/group. **m:** The probability of making a turn in motion direction was the same for all fish before any group was subjected to a sleep disruption protocol (mixed ANOVA: $p_{grp} = 0.67$, F(1,62) = 0.178; Tukey's HSD: $p_{0\%} = 0.82$, $p_{25\%} = 0.89$, $p_{50\%} = 0.71$, $p_{100\%} = 0.52$, n = 32/group). **n:** The bout rate between both undisturbed groups of fish was the same (mixed ANOVA: $p_{grp} = 0.75$, F(1,62) = 0.1; Tukey's HSD: $p_{0\%} = 0.54$, $p_{25\%} = 0.68$, $p_{50\%} = 0.72$, $p_{100\%} = 0.82$, n = 32/group). **o:** No difference in reaction time was observed. The inset shows the mean time to the first bout (independent samples t-test: p = 0.23, t = 1.2, df = 62, n = 32/group). **p:** Turning angle distribution was similar between groups. Graphs represent mean ±SEM with the responses of individual fish overlaid in circles.

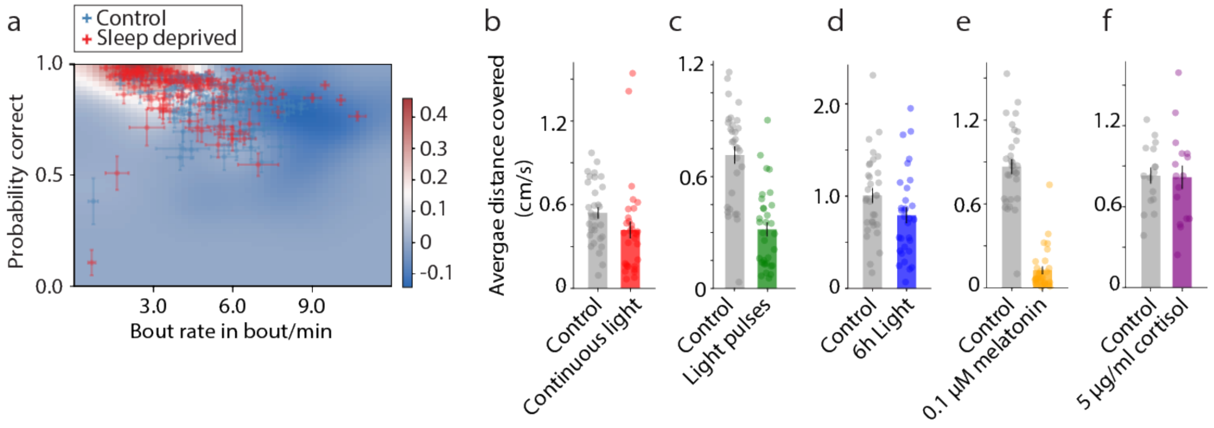


**Figure S3: A decrease in bout rate is correlated to increased correctness and reduced exploration in the OMR task. a:** The uniqueness map displays the difference in density (uniqueness) between disrupted (light, light pulses and 6h light) and sibling controls. Colors indicate the extent of uniqueness with

0 meaning full overlap in density between both groups. Sleep-disrupted fish dominate red areas while control fish dominate dark blue areas. The overlaying scatterplot ±SEM shows the bout rates and the probabilities for correct turns per fish. A negative correlation was found between bout rate and performance for both sleep-disrupted (Spearman's rank correlation coefficient: $\rho = -0.36$, $p = 4.1^{-5}$, $n = 48$) and control animals (Spearman's rank correlation coefficient: $\rho = -0.43$, $p = 3.9^{-7}$, $n = 48$). **b:** Sleep disruption with continuous light slightly reduced exploration measured as average distance covered during the OMR task (independent samples t-test: $p = 0.91$, $t = 1.72$, $df = 61$, $n = 32$/group). **c:** Sleep disruption with light pulses decreased exploration (independent samples t-test: $p < 0.001$, $t = 6.68$, $df = 61$, $n = 32$/group). **d:** Sleep disruption with 6 h of premature light slightly reduced exploration (independent samples t-test: $p = 0.06$, $t = 1.86$, $df = 61$, $n = 32$/group). **e:** Exploration was reduced post-melatonin administration (paired t-test: $p < 0.001$, $t = 16.16$, $df = 30$, $n = 32$). **f:** Cortisol administration did not alter spontaneous exploration measured as the average distance covered (paired t-test: $p = 0.82$, $t = 0.24$, $df = 15$, $n = 16$). Bar graphs represent mean ± SEM with the responses of individual fish overlaid in circles.

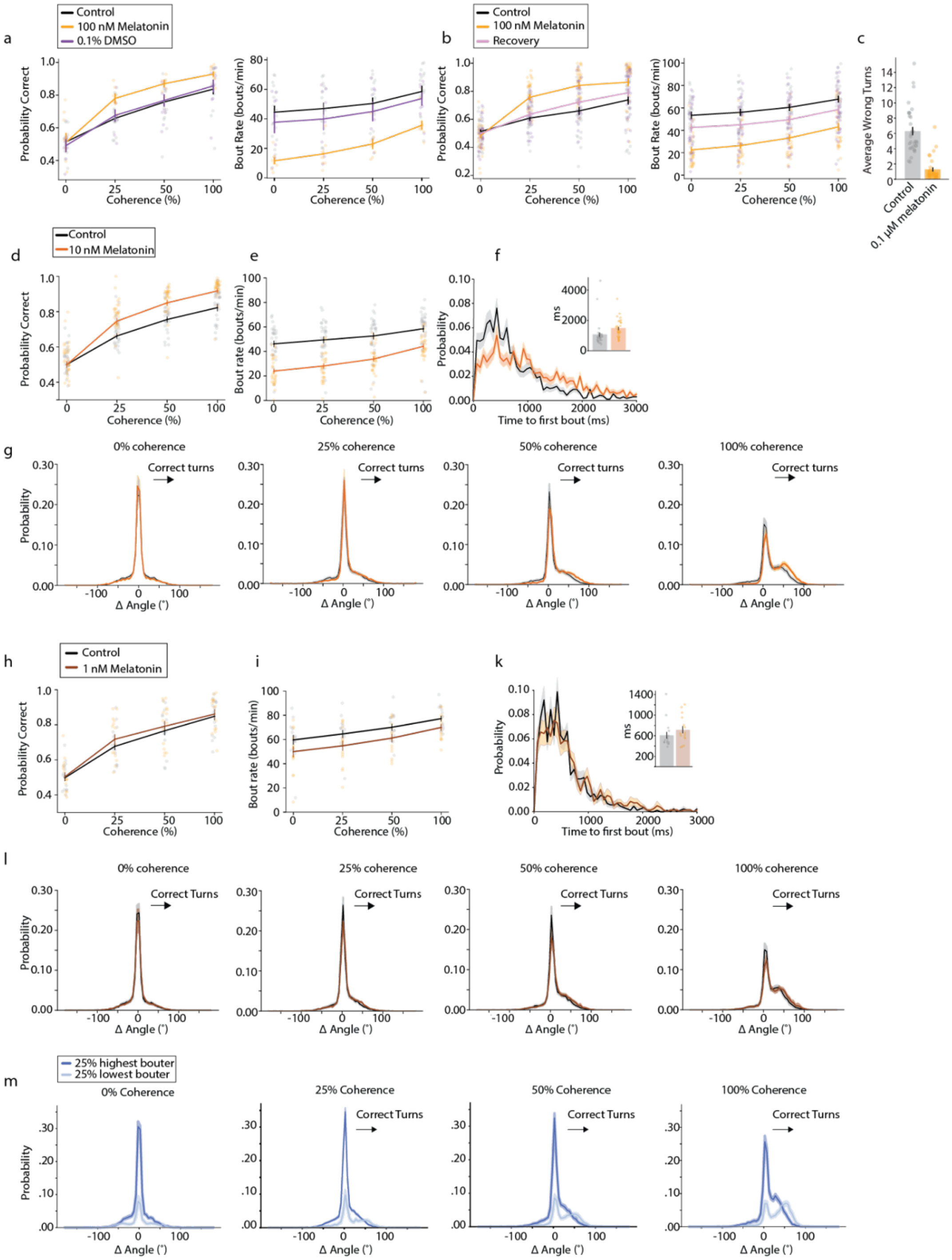


**Figure S4: Melatonin reversibly increases reaction time and performance in a concentration-dependent manner. a:** To exclude any effects of the DMSO solvent on behavior, the optomotor response was tested before treatment and after treatment with either 100 nM melatonin or 0.1% DMSO. Left: Melatonin-treated fish performed better than control and DMSO-treated groups (repeated measures ANOVA: p + 0.047, F (2,14) = 3.83; Tukey's HSD *control vs melatonin*: $p_{0\%}$ = 0.9, $p_{25\%}$ = 0.002, $p_{50\%}$ = 0.029, $p_{100\%}$ = 0.12; Tukey's HSD *control vs DMSO*: $p_{0\%}$ = 0.73, $p_{25\%}$ = 0.83, $p_{50\%}$ = 0.9, $p_{100\%}$ = 0.9, $n_{ctrl}$ = 16, $n_{mel}$ = 8, $n_{DMSO}$ = 8). Right: Bout rate was decreased for melatonin-treated fish compared to the control, but not for fish treated with DMSO (repeated measures ANOVA: p < 0.001, F (2,14) = 14.17; Tukey's HSD *control vs melatonin*: $p_{0\%}$ < 0.001, $p_{25\%}$ < 0.001, $p_{50\%}$ < 0.001, $p_{100\%}$ = 0.002; Tukey's HSD *control vs DMSO*: $p_{0\%}$ =

0.6, $p_{25\%}$ = 0.58, $p_{50\%}$ = 0.69, $p_{100\%}$ = 0.69, $n_{ctrl}$ = 16, $n_{mel}$ = 8, $n_{DMSO}$ = 8) **b:** Optomotor response tested in larvae before, during, and after treatment with 100 nM melatonin. n = 32/group. Left: The correctness recovered to levels seen before melatonin treatments within 20 minutes (repeated measures ANOVA: p < 0.001, F (2,60) = 26.07; Tukey's HSD *control vs melatonin*: $p_{0\%}$ = 0.089, $p_{25\%}$ < 0.001, $p_{50\%}$ < 0.001, $p_{100\%}$ = 0.006; Tukey's HSD *control vs recovery*: $p_{0\%}$ = 0.47, $p_{25\%}$ = 0.53, $p_{50\%}$ = 0.133, $p_{100\%}$ = 0.42, n = 32). Right: The bout rate recovered to levels similar to before melatonin treatment within 20 minutes (repeated measures ANOVA: p < 0.001, F (2,62) = 44.54; Tukey's HSD *control vs melatonin*: $p_{0\%}$ < 0.001, $p_{25\%}$ < 0.001, $p_{50\%}$ < 0.001, $p_{100\%}$ < 0.001; Tukey's HSD *control vs recovery*: $p_{0\%}$ = 0.04, $p_{25\%}$ = 0.04, $p_{50\%}$ = 0.051, $p_{100\%}$ = 0.14, n = 32). . **c:** The total number of wrong turns over all coherences was decreased post administration with 100 nM melatonin (paired t-test: p < 0.001, t = 8.27, df = 30, n = 32). **d - g:** Fish treated with 10 nM melatonin showed improved performance and reaction time in the visual decision-making task. n=32/group. **d:** Melatonin-treated fish showed improved performance compared to before the treatment (repeated measures ANOVA: p < 0.001, F (1,30) = 19.77; Tukey's HSD: $p_{0\%}$ = 0.9, $p_{25\%}$ < 0.001, $p_{50\%}$ < 0.001, $p_{100\%}$ < 0.001, n = 32). **e:** Melatonin-treated fish had a reduced bout rate (repeated measures ANOVA: p < 0.001, F(1,30) = 48.53; Tukey's HSD: $p_{0\%}$ < 0.001, $p_{25\%}$ < 0.001, $p_{50\%}$ < 0.001, $p_{100\%}$ < 0.001, n = 32). **f:** Melatonin treatment increased the larvae's reaction time after the onset of the 100% coherent stimulus (paired samples t-test: p = 0.026, t = -2.34, df = 30, n = 32). The inset shows the mean time to the the first bout. **g:** Treated fish had an increased turning angle while making more correct turns. Their spontaneous swims were not compromised (0% coherence). **h - l:** Fish treated with 1 nM melatonin did not show improvement. n = 16. **h:** The probability correct was marginally increased in fish after treatment (repeated measures ANOVA: p = 0.28, F (1,14) = 1.25; Tukey's HSD: $p_{0\%}$ = 0.64, $p_{25\%}$ = 0.15, $p_{50\%}$ = 0.47, $p_{100\%}$ = 0.72, n = 16). **i:** Treatment with 1 nM melatonin decreased the bout rate marginally (repeated measures ANOVA: p = 0.015, F (1,14) = 7.68; Tukey's HSD: $p_{0\%}$ = 0.12, $p_{25\%}$ = 0.075, $p_{50\%}$ = 0.058, $p_{100\%}$ = 0.051, n = 16)**. k:** The reaction time was minimally increased after treatment (paired samples t-test: p = 0.12, t = -1.67, df = 14, n = 16). The inset shows the mean time to the first bout. **l:** Turning angles were slightly shifted towards bigger turns in treated fish. **m:** Low-bouting fish showed decreased probabilities for forward swims and turns for all coherences, but a slight shift to bigger turning angles, especially for a 100% coherent stimulus. Graphs represent mean ±SEM with the responses of individual fish overlaid in circles.

#### Computational model

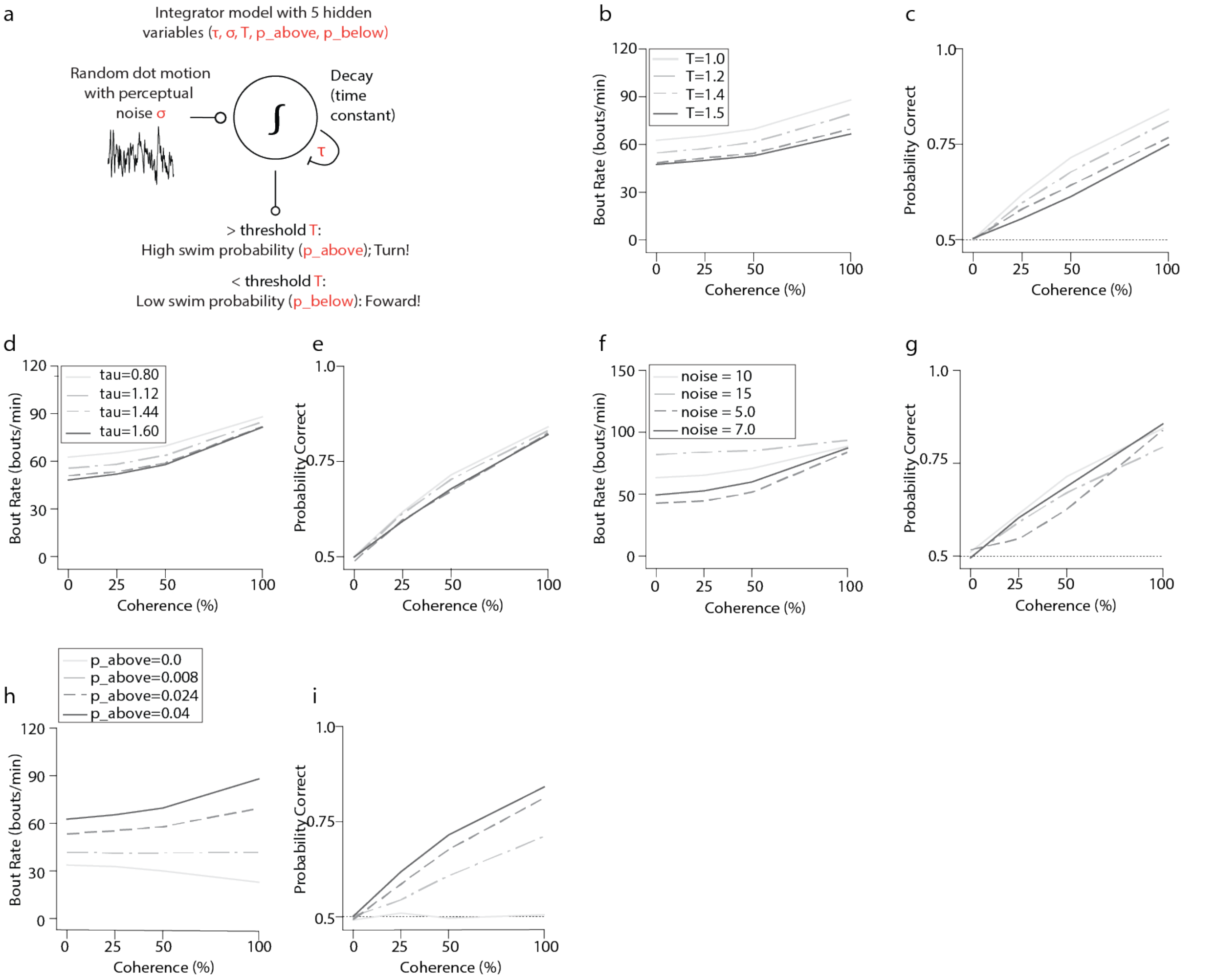


**Figure S5. Varying parameters of the drift diffusion model. a:** Integrator model. Figure adapted from Harpaz et al. (2021) [33]. Increasing the dynamic threshold decreased the bout rate (**b**) but diminished performance (**c**). Increasing tau decreased the bout rate (**d**) but did not affect the performance (**e**). Varying the perceptual noise, altered the bout rate (**f**) but decreased the performance (**g**). Decreasing the bouting probability for turns ($p_{above}$) (**h**) also decreased the performance (**i**).

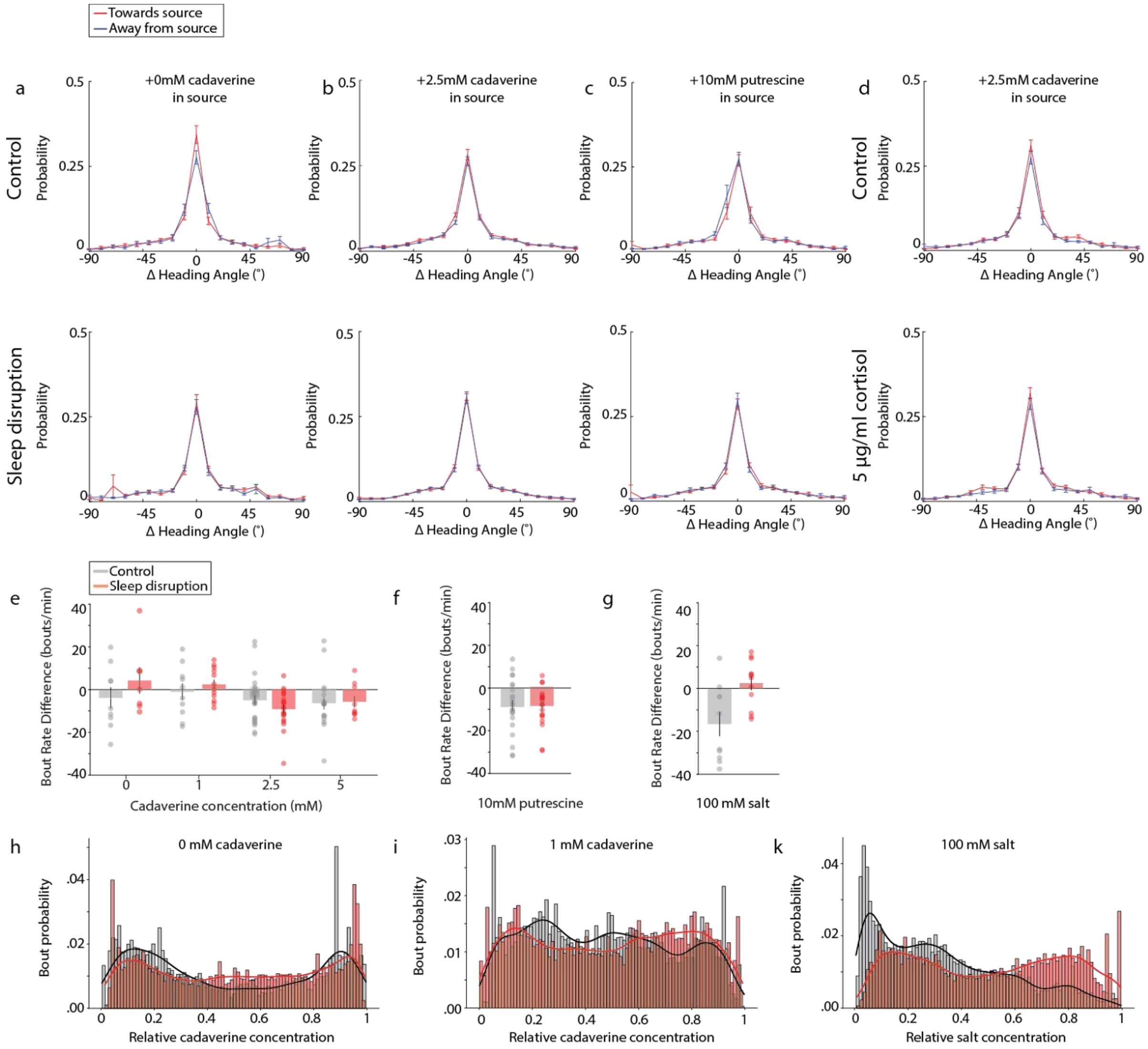


**Figure S6. Turning angles do not change between conditions. a:** Without the presence of an odor neither control nor sleep-disrupted fish showed a difference in turning angle distributions between either side of the setup. **b**: Control and sleep-disrupted fish did not show a change in turning angle distribution in response to 2.5 mM cadaverine. **c:** Control and sleep-disrupted fish did not show a change in turning angle distribution in response to putrescine. **d:** Larvae did not show a change in turning angle distributions before or after cortisol treatment. **e:** Sleep-disrupted fish did not change their bout rates close to the cadaverine source (Wilcoxon rank sum test: 0mM: p = 0.27, $n_{ctrl}$ = 10, $n_{SD}$ = 10; 1mM: p = 0.39, $n_{ctrl}$ = 10, $n_{SD}$ = 13; 2.5mM: p = 0.32, $n_{ctrl}$ = 27, $n_{SD}$ = 22; 5mM: p =0.68, $n_{ctrll}$ = 19, $n_{SD}$ = 14). **f:** Sleep-disrupted fish likewise did not change their bout rate close to the putrescine source (Wilcoxon rank sum test: p = 0.95, $n_{ctrl}$ = 33, $n_{SD}$ = 30). **g:** Sleep-disrupted fish increased their bout rate in response to salt (Wilcoxon rank sum test: p = 0.01, $n_{ctrl}$ = 10, $n_{SD}$ = 16). **h-k:** Normalized bout probability across relative odor concentrations. **h:** In the absence of an odor, the bout probabilities were evenly distributed for well-rested and sleep-disrupted fish, with a slight increase towards the relative odor concentrations of 0 and 1. **i:** For 1 mM cadaverine, both control and sleep-disrupted groups showed a decreased bout count in high cadaverine concentrations. **k:** Sleep-disrupted fish did not return earlier when encountering salt, but showed an increased bout count in high salt areas compared to controls. Graphs represent mean ±SEM. Histograms show the normalized bout count probabilities across 85 bins including a kernel density estimate.

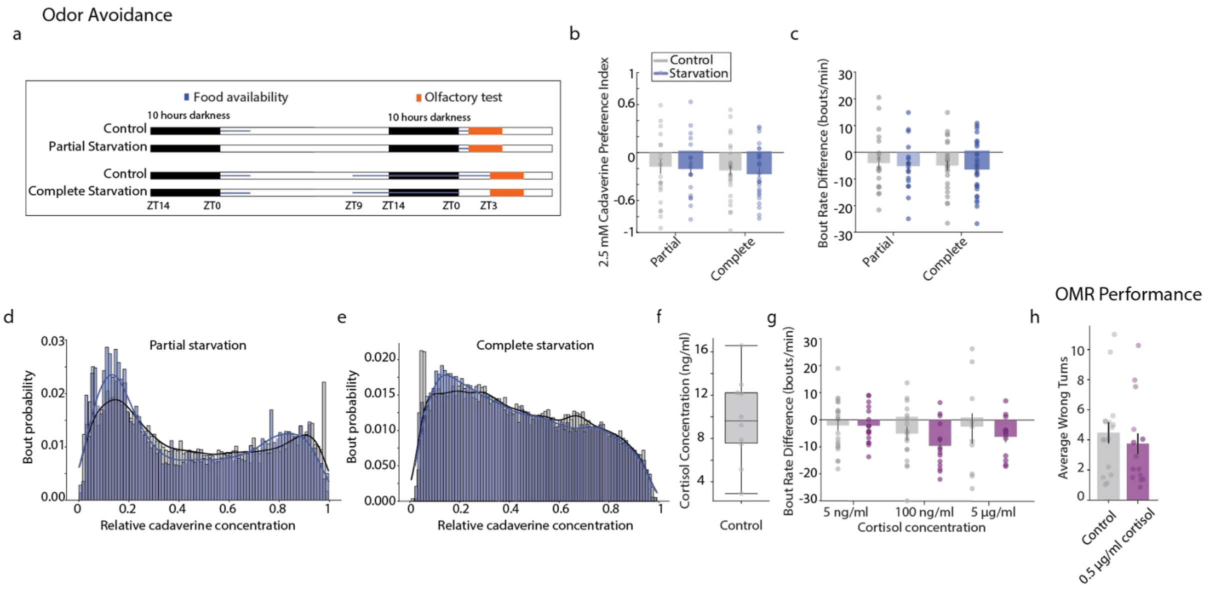

**Figure S7. Food deprivation cannot mimic behavioral changes of sleep disruption. a:** Fish were starved with two different schedules. For *partial starvation* control fish were fed according to the regular schedule at ZT0 every morning. Partially starved fish were not fed at ZT0 the day before experimentation, however, both control and partially starved fish were fed at ZT0 immediately preceding experimentation. For *complete starvation*, control and starved fish were fed at ZT0 the day before experimentation and again at around ZT9 the night before experimentation. Control fish were again fed at ZT0 right before the experiment; however, the starved fish were not. **b:** No significant changes in the avoidance behavior were seen between the control and either group of starved fish (*partial starve*: Wilcoxon rank sum test: $p = 0.90$, $n_{ctrl} = 22$, $n_{starvel} = 18$; complete starve: Wilcoxon rank sum test: $p = 0.78$, $n_{ctrl} = 27$, $n_{starve} = 28$). **c:** Neither partial nor complete starvation changed the fish's bout rates (Wilcoxon rank sum test: partial starve: $p = 0.99$, $n_{ctrl} = 22$, $n_{starve} = 18$; complete starve: $p = 0.64$, $n_{ctrl} = 27$, $n_{starve} = 28$ ). **d:** Partial starvation did not discernibly alter the bout distribution along the cadaverine gradient compared to unstarved controls. **e:** Complete starvation did not discernibly alter the bout distribution along the cadaverine gradient compared to unstarved controls. **f:** Cortisol concentration of all control groups. n=9. Each circle represents a group of 30 fish. **g:** Cortisol-treatment did not alter the bout rate difference (Wilcoxon rank sum test: 5ng/ml cortisol: $p = 0.85$, $n_{ctrl} = 27$, $n_{cort} = 21$; *100 ng/ml cortisol*: $p = 0.23$, $n_{ctrl} = 19$, $n_{cort} = 15$; *5 µg/ml cortisol:* $p = 0.90$, $n_{ctrl} = 11$, $n_{cort} = 16$). **h:** The number of wrong turns in the OMR task did not discernibly change after cortisol administration (paired t-test: $p = 0.16$, $t = 1.47$, $df = 15$, $n = 16$). Bar graphs represent mean ± SEM with the responses of individual fish overlaid in circles unless stated otherwise. Histograms show the normalized bout count probabilities across 85 bins including a kernel density estimate.
